# Building the Mega Single Cell Transcriptome Ocular Meta-Atlas

**DOI:** 10.1101/2021.03.26.437190

**Authors:** Vinay S Swamy, Temesgen D Fufa, Robert B Hufnagel, David M McGaughey

## Abstract

The development of highly scalable single cell transcriptome technology has resulted in the creation of thousands of datasets, over 30 in the retina alone. Analyzing the transcriptomes between different projects is highly desirable as this would allow for better assessment of which biological effects are consistent across independent studies. However it is difficult to compare and contrast data across different projects as there are substantial batch effects from computational processing, single cell technology utilized, and the natural biological variation. While many single cell transcriptome specific batch correction methods purport to remove the technical noise it is difficult to ascertain which method functions works best. We developed a lightweight R package (scPOP) that brings in batch integration methods and uses a simple heuristic to balance batch merging and celltype/cluster purity. We use this package along with a Snakefile based workflow system to demonstrate how to optimally merge 766,615 cells from 33 retina datsets and three species to create a massive ocular single cell transcriptome meta-atlas. This provides a model how to efficiently create meta-atlases for tissues and cells of interest.

## Introduction

### A plethora of single-cell transcriptome studies in the retina

The retina contains a multitude of cell types that, in total, are responsible for turning light information into signal for the brain to interpret as vision. Very briefly, the photoreceptors (rods and cones) are responsible for capturing the photons. The retinal pigmented epithelium (RPE) behind the photoreceptors physically support the rods and cones by processing byproducts of the visual cycle. Müller glia serve as support cells for the neurons. The retinal bipolar cells transmit the electrical signal from the photoreceptors to the retinal ganglion cells. Horizontal and amacrine cells regulate and help interpret signals from the photoreceptors. The signal is relayed via the retinal ganglion projections through the optic stalk to the brain (for review see).^1^ Since 2000 many groups have investigated gene expression in small numbers of individual cells of the retina.^2–5^

The recent introduction of lower cost and high throughput single cell sequencing technology has led to an explosion of research across many fields. As of early 2021, over 40 million cells have been sequenced across over 1,200 studies and the average size of each study starting in 2020 is over 100,000 cells.^6^ The retina was used as the source tissue in one of the earliest works in the high throughput single cell transcriptomics field.^7^ As of late 2020, over twenty published studies, cumulatively containing over a million cells, have used single cell technology to profile cell type specific gene expression patterns, cell fate trajectory, tissue and cell differentiation, and disease perturbation across multiple mammalian species.^7–15, 15–19, 19–29^

While the gene - cell count tables are generally made available in repositories like the Gene Expression Omnibus (GEO), there are no requirements to uniformly process the data. This means the count tables cannot be used in cross-study comparisons as even small differences in the computational pipeline (aligner, transcriptome reference, etc.) create study-specific effects. This issue can be addressed only by re-quantifying the data in a uniform computational environment. Fortunately, due to the continued development of computationally light-weight gene quantification tools in the single-cell space (e.g kallisto bustools, alevin-fry), re-quantification does not require massive compute and time resources.^30, 31^

Still, even after re-quantification under identical computational conditions there remain study specific batch effects due to the diversity in single cell technologies used and variation in tissue handling and processing across each scientific group. The single cell community has recognized that removal of these technical (also referred to as batch) effects is a critical issue and have independently developed many tools, though it remains unclear which tools and parameters are optimal for a particular dataset.^32–43^ Reinforcing this point, a couple groups have quantified performance of multiple methods in a consistent framework across several test datasets, finding no consistently best approach.^44, 45^

### The projectable meta-atlas

We propose that by re-processing publicly available raw single cell transcriptome data in a consistent bioinformatic framework and optimally using batch correction tools we can create a meta-atlas of retina single cell transcriptomes. As there are thousands of possible permutations of single cell tools, references, and parameter choices, we create our meta-atlas (which we refer to as the single cell Eye in a Disk or scEiaD) by benchmarking integration outcomes across multiple important single cell RNA-seq processing parameters (batch removal method, number of hyper-variable genes (HVGs), clustering resolution, etc.). The benchmarking system we developed uses a wide variety of metrics that combine in the R package scPOP (single-cell Pick Optimal Parameters). The scEiaD will be of utility to two communities. First, the ocular community who can both search scEiaD for gene expression across many dimensions (e.g. cluster, cell type, study) and project their own single cell data onto scEiaD for comparison and rich automatic cell labeling. Second, the computational community can use this very large, well-curated dataset to test algorithms for compute efficiency and performance in a diverse environment. As we believe data re-use is a powerful and efficient approach to facilitate discovery, we provide our meta-atlas code-base, the meta-atlas in several data formats, and propose general guidelines to optimally create custom meta-atlases.

## Results

### We identify 33 ocular scRNA datasets across 3 species

The first step in building a meta-atlas is identifying studies to draw the data from. We identified ocular single cell RNA sequencing (scRNA) studies by querying PubMed, the Sequence Read Archive (SRA), and the European Nucleotide Archive (ENA) for the inclusive terms “retina,” “single cell,” “scRNA,” “ocular,” “eye,” “transcriptome.” We then hand filtered the results to only keep ocular and normal (non-perturbed or mutagenized) data from single cell RNA-seq technology. On December 2020 we identified 33 deposited datasets that have been published in 27 publications (Figure 1). To provide a non-ocular reference we also downloaded the raw sequence data from the Tabula Muris project for re-processing. In cases where the fastq file from from the SRA was not processed properly (always 10x v2 or v3), we acquired the 10x bam files (via SRA or personal correspondence) and re-extracted the fastq. After downloading all the data we had 11 TB across 2470 fastq file sets.

**Figure 1:**
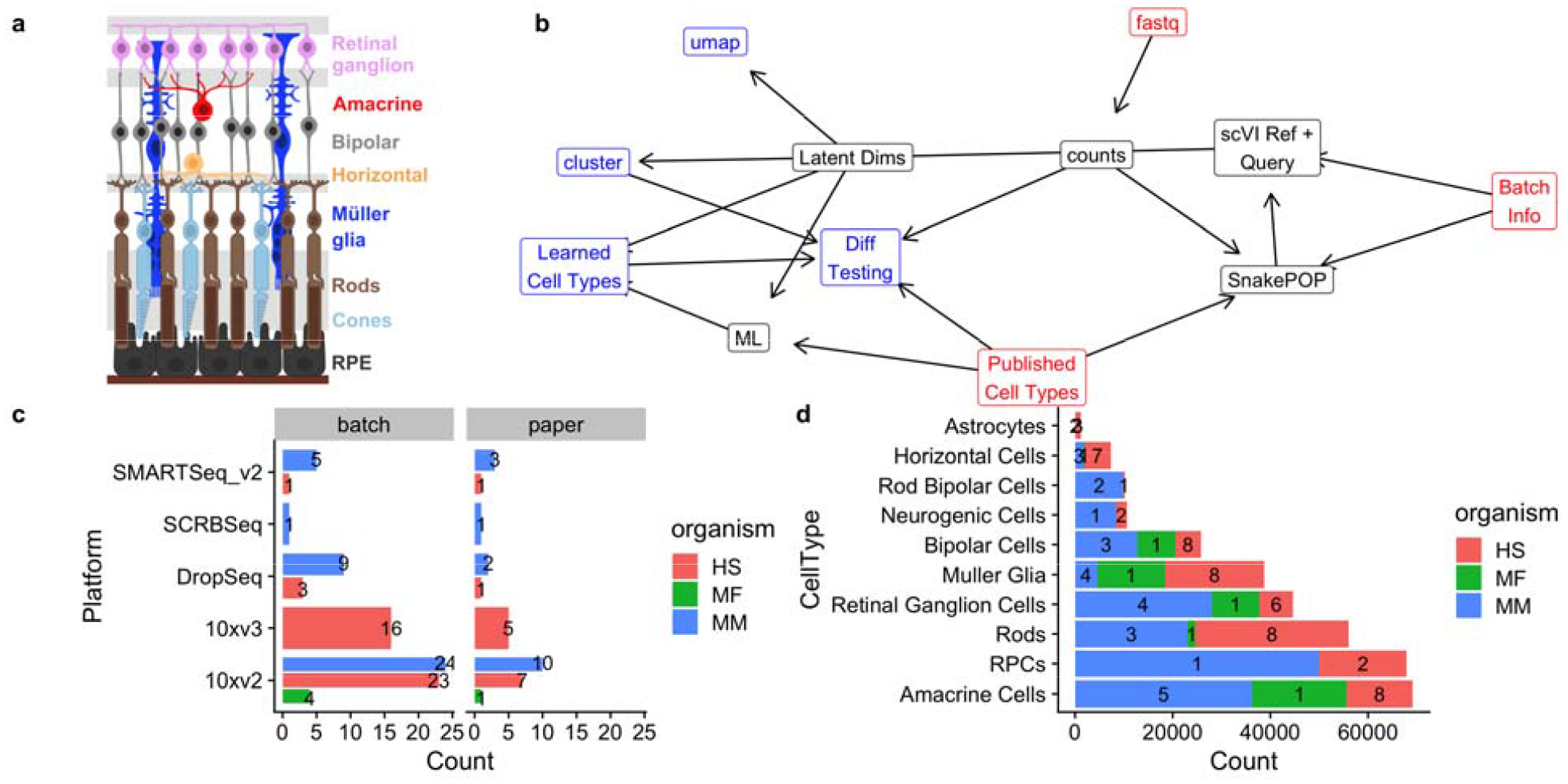
a. Schematic of the retina with major cell types delineated b. Simplified directed workflow of major steps in scEiaD creation from raw counts to gene counts, benchmarking optimal integration methods (SnakePOP) to produce batch corrected latent dimensions (Latent Dims), then downstream analysis outputs like clustering, differential gene testing (Diff Testing), and 2D UMAP visualization. c. Counts of published papers and batches (unique biological samples) for each scRNA technlogy, split by organism d. Cell type counts extracted from published studies for the more common retina cell types, split by species. Count of study accessions for each species overlaid on bar plot.

### Transcriptome quantification across multiple technologies

Droplet and well based scRNA-seq technologies require different quantification approaches as the former have UMI and multiple cells are quantified within a single file. We wrote a Snakemake based pipeline (SnakeQUANT, see methods) to quantify and merge both droplet and well based technologies into a single matri for downstream processing. For well based data we perform both gene and isoform level quantification; for droplet base technologies we quantify both exonic and intronic gene-level expression to facilitate calculation of RNA velocity. In total we quantify 6.7e+10 molecules, finding 1.1e+10 unique molecules with a mean pseudoalignment rate of 66.5%. Across the 766,615 cells (post QC) we have an average of 3,410 RNA counts across 978 unique genes (see Supplemental File kallisto_stats.tsv and splicing_stats.tsv for more details).

### 1,204,269 cells before quality control

Gene-level counts were quantified with the kallisto bustools pseudo-aligner for both the droplet and well based samples. After empty droplet removal, we had 1,204,269 cells. We then removed cells which had more than 10% mitochondrial reads across all gene counts, fewer than 200 unique genes quantified, or were identified as an *in silico* doublet (see methods). After these quality control (for review see^46^) steps we were left with 766,615 cells (Supplemental Figure 1).

A core objective of many scRNA based studies is labeling the cell types. As this information is crucial to assess dataset integration and provide an accurate reference for user querying, we extracted individual cell labels with a combination of inspecting the GEO web site, supplemental information from the publication, web resources (e.g. a web app was created for the paper), and personal correspondence. After normalizing cell type name nomenclature, we obtained labels for 375,966 cells across 33 cell types (Supplemental Table 1).

### Running 11 tools in a Snakemake-based system

Disentangling the technical and biological effects when integrating multiple datasets is crucial. We define batch as each unique biological sample and assume each study is at least one unique sample. We studied the metadata and methods of each study to identify the unique biological samples. Within the current scEiaD data set we identified 86 batches across 33 deposited datasets in 26 published papers.

A wide variety of methods have been written for scRNA-seq integration. As we were uncertain which would perform the best, we ran 11 tools with a commonly used set of key parameters like number of highly variable genes (HVG), number of latent dimensional space, and the number of nearest neighbors for the louvain clustering algorithm. The Snakemake system was used to automate the running of the wide variety of tools. In total 5,591 jobs were run to quantify gene expression and build the unified Seurat objects (SnakeQUANT) and 2,446 jobs were run to assess integration performance (SnakePOP).

The two key metrics which have to be balanced in order to optimize integration performance are cell type or cluster purity (where different cell types or clusters should be homogenous) and batch mixing (the same cell types should be similar across independent studies). While these can be visually assessed by looking at marker gene expression across the 2D UMAP projection, it is more rigorous and scalable to quantify these diametrically opposed characteristics.

### scPOP wraps several different methods for measuring integration performance

Multiple methods have been proposed to quantitatively evaluate batch correction. Some of these metrics evaluate the concordance between sets of labels, while other compute distances between the individual data points of a given set of labels. While any one of these methods can be useful, we propose that calculating and evaluating them in tandem provides greater accuracy for dataset integration. We developed the R package scPOP, a lightweight, low dependency R package which brings together the Local Simpson Index (LISI), Average Silhouette Width (ASW), Adjusted Rand Index (ARI) and Normalized Mutual Information (NMI) metrics from the R packages Harmony, kBET and aricode, respectively. The LISI and ASW were used to measure batch mixing (where lower is better), cell type mixing (higher is better), and cluster mixing (higher is better). NMI and ARI were used to assess the consistency of cell type to cluster assignment (where 1 is perfect correspondence between cluster and cell type).

To visualize the interplay between batch mixing and cell type distinction we plot the batch mixing LISI score (which has been multiplied by −1) on the y-axis (higher is better) against the cluster LISI on the x-axis (higher is better). The best performer on both metrics will be in the top right corner (Supplemental Figure 2a). In the same manner we plot the silhouette metric (Supplemental Figure 2 b). To merge the different scores we define *sumZScale* = ∑*scale*(*m* * *Weight*) where m is a metric (LISI by batch, LISI by cluster, LISI by cell type, silhouette by batch, silhouette by cluster, silhouette by celltype, NMI, and ARI). Weight is set to one, but can be either explicitly set or randomly chosen should it be desired to change the influence of certain metrics.

On one extreme we have ComBat, which merges together different batches very well, but also mixes together the distinct cell types (Figure 2a). The other extreme is not using any batch integration method, where you see very distinct groups of cells, but also each nearly study is has a distinct region in the UMAP (Figure 2b). In our scEiaD dataset we see that methods like ComBat, Harmony, scArches, CCA are weighed more towards batch mixing then cluster and cell type purity. Scanorama and bbknn prioritize cleanly separating the clusters. With our scEiaD meta-atlas insct, trVAE, and magic do not perform particularly well in batch mixing or cluster purity. Overall, CCA, fastMNN, and scVI perform best on this particular dataset. For further qualitative investigation of the integration performance we provide the UMAP visualizations of each integration method, normalization, and latent dimensions colored by cell type, study accession, or organism as supplementary files.

**Figure 2:**
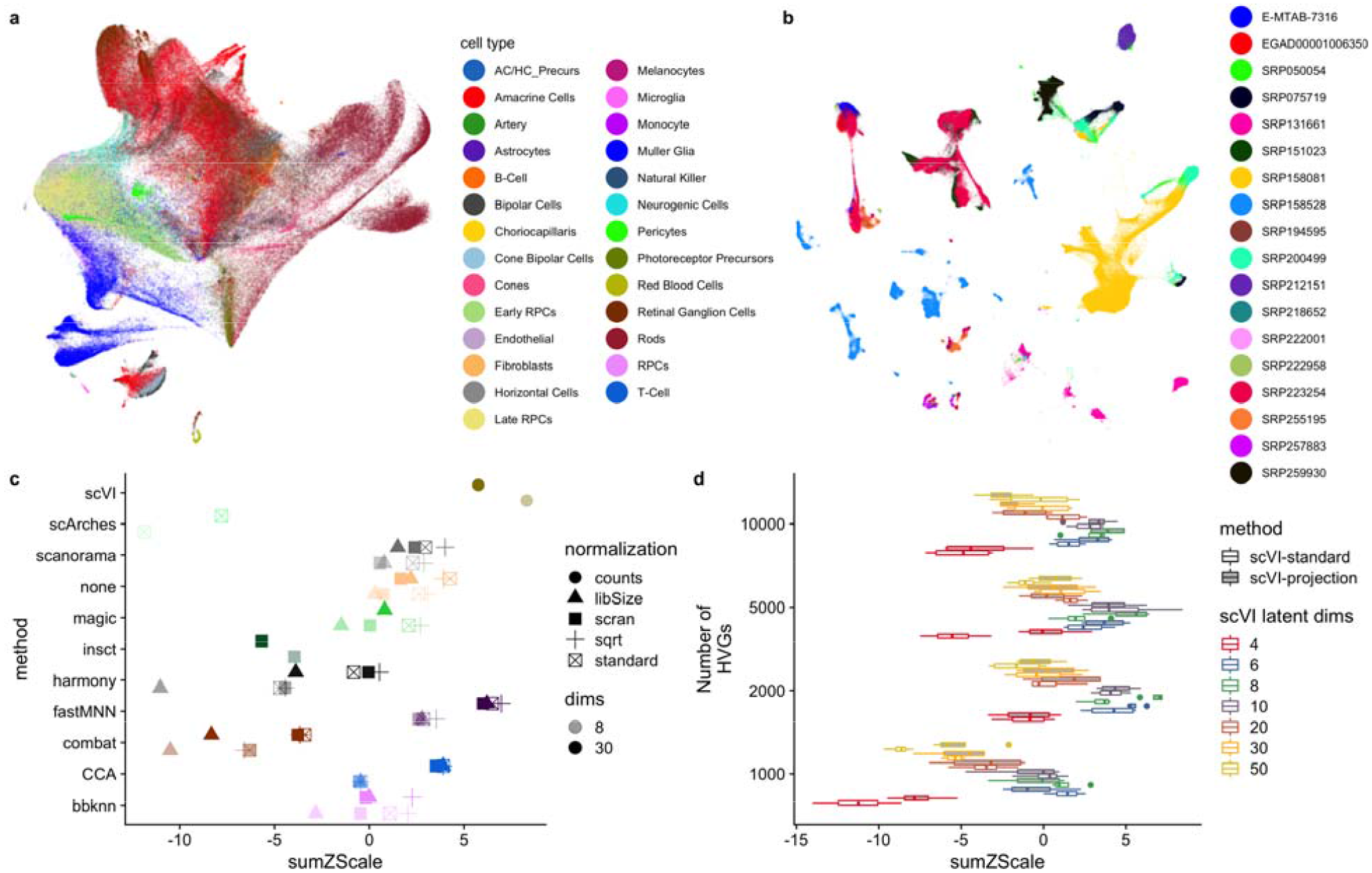
a. Example of a method (combat) which has a high level of batch blending, but poor separation of cell types (colored by cell type). b. no batch correction cleanly separates cell types but does not mix batches (colored by study). c. sumZScale (higher is better) for each method across a variety of data normalizations. All methods shown here use 2000 HVG, louvain clustering with knn 20, and 8 or 30 latent dimensions. Each color is a different method. d. Boxplot of 4 different clustering resolutions for across 1000 to 10000 HVG numbers and 4 to 50 scVI latent dimensions. Open boxes are using scVI-standard and gray boxes are scVI-projection (human reference with the remaining data projected)

As the LISI and silhouette metrics provide independent cluster and cell type (“purity”) and batch (“mixing”) scores we looked to see whether the sumZScale scoring is highly influenced by changing the weight placed on purity or mixing by multiplying either by a multiplier of three (Supplemental Figure 3). While scVI still performed well, no matter the weight chosen, there were some changes when more weight was placed on batch mixing. Most notably fastMNN with 8 latent dims performed better than fastMNN with 30 latent dims. In a reveral of the fastMNN results, CCA with 30 latent dimensions performed better than CCA with 8 latent dimensions. We also bootstrap random weights across each individual metric and get similar results (Supplemental Figure 4). Across the 11 integration methods we tested, scVI achieved the highest sumZScale for our scEiaD meta-atlas and furthermore is a desirable choice because of its very short run time (scVI can complete in less than an hour while fastMNN takes over 12 hours and CCA takes more than 24 hours).

### Different normalization methods alter integration performance

There are several normalization approaches that have been used or published. The “standard” approach that the popular analysis packages Seurat and scanpy use by default is to, per cell, divide the counts by the sum counts for the cell, multiply by a scaling factor, then log transform. This helps make the count distribution more normal, which is an assumption that many algorithms require. In contrast, the scran normalization method groups cells into pools and normalizes across the pool summed counts instead of the individual cell counts. We also use the square root (sqrt) normalization which replaces the log transformation with a square root. Library size (libSize) normalization omits the sqrt or log transformation. Finally some methods, like scVI, directly use the raw counts for modeling the data.

As expected the libSize normalization which omits the log or square root scaling generally performs the worst (Figure 2c). We see that the remaining normalization techniques alter the batch correction performance, though the outcome differs across the different methods. We also see that changing the number of latent dimensions (8 or 30) can occasionally dramatically change performance. These results demonstrate the importance of assessing performance in a rigorous manner across many parameters.

### Further optimization of scVI with grid search and projection

To find the best set of parameters for scVI in out dataset we did a grid search across key parameters: HVG, latent dimensions, and k nearest neighbors. Furthermore we used a recent advance in scVI capability (>= versio 0.8.0) adapted from scArches that allows one to build a reference model and query or project the new cells onto it.^40^ We refer to this projectable model as “scVI-projection,” and the previous scVI model as “scVI-standard.” We built a scVI-projected model trained on human cells and then projected the mouse and macaque data onto it. We the compared scVI-projection against the against the previous scVI-standard across all the previously mentioned parameters. Using scPOP we first saw that the scVIprojection approach generally performed better than running scVI with all of the data (Figure 2d). We found the optimal overall parameters to be 5000 HVG, 8 latent dimensions, and with 5 k-nearest neighbors for the cluster finding. We also varied the UMAP projection values of nearest neighbors and minimum distance to qualitatively pick a 2D projection, selecting a minimum distance of 0.1 and 50 nearest neighbors.

### High accuracy xgboost ML model built to label unknown cell types across all technologies

To further study cell type specific expression patterning we needed to label the 414,106 unlabeled cells. Traditionally this is done by clustering the cells, then using cell type specific markers to label the clusters. However, as we had hundreds of thousands of expert labeled cells across 17 publications (Figure 1a) we built a xgboost-based machine learning model that used 2/3 of the labeled cells as a training set (see methods for more details) to train a cell type predictor for scEiaD input. The trained model was used to predict the cell type assignments for all cells in scEiaD. In this manner we both label most cells (cells which cannot assign a cell type with a probability above 0.5 were left unlabeled) and correct a small number of probable mislabels in the truth set (Figure 3a).

**Figure 3:**
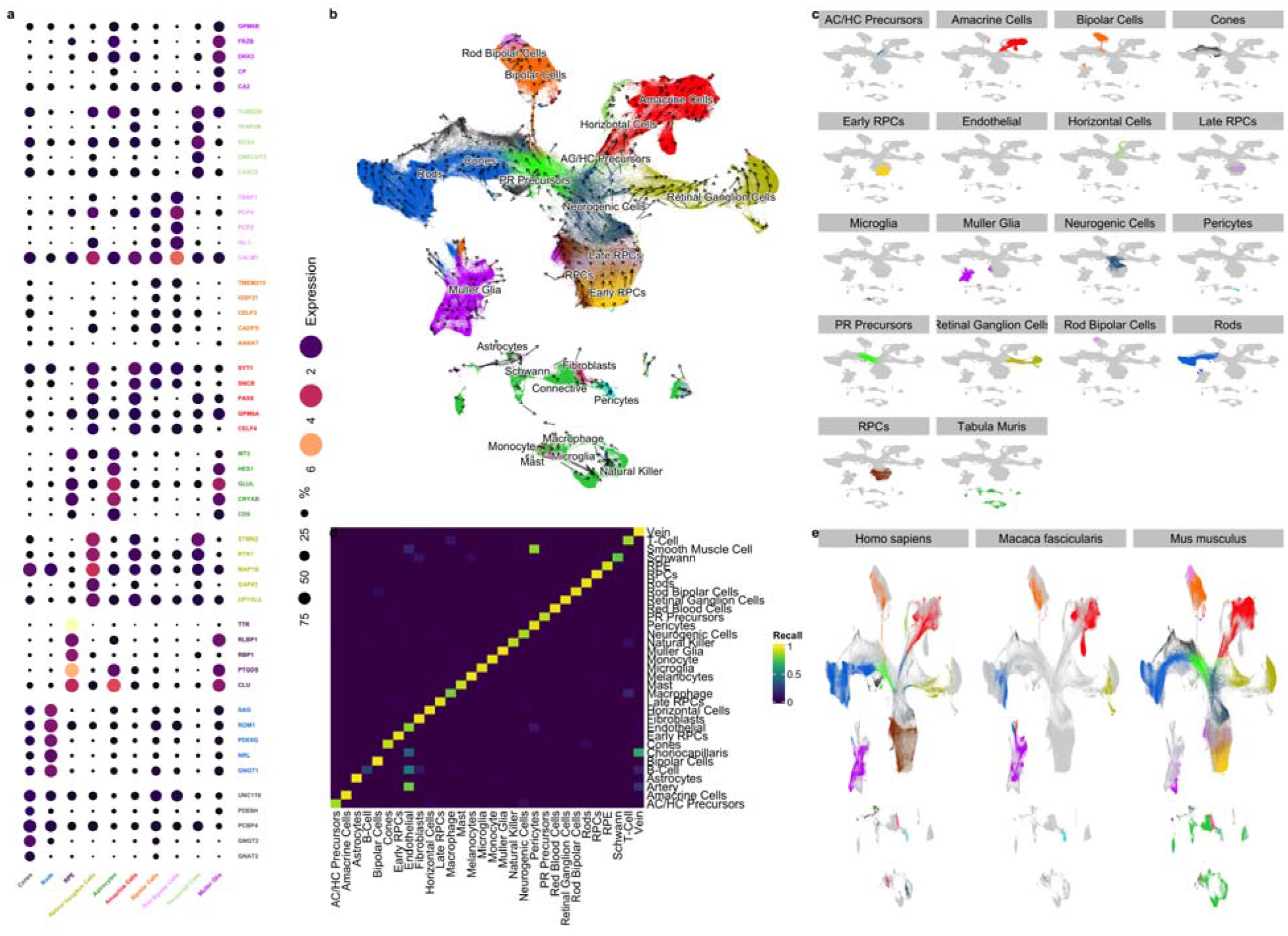
a. Top genes that are differentially expressed across the major cell types of the retina (PR is short for Photoreceptor). Genes are colored by which cell type they are differentially expressed in. The dot size is proportional to the percentage of those cells that have detectable levels of the gene. The color of the dot is the log2 scaled CPM expression. b. 2D UMAP projection of scEiaD, colored by cell type (Tabula Muris data is gray). Arrows are scvelo RNA velocity. Longer arrows are cells with higher velocity (relatively more unspliced transcripts).c. Facet plot that demonstrates how each major cell type of the retina is contained within a distinct space. d. Confusion matrix of cell type prediction performance of our xgboost labeller between predicted (x axis) and known (y axis) using data withheld from the machine learner. Most of the cell types are indeed labeled as their true type. e. Faceting of 2D UMAP by species and colored by cell type demonstrate how the major cell types of the retina share space with like cell types, despite being from mouse, human, and macaque.

As a brief case study of our ML performance, we look at the cell type labels assigned to the Shekhar et al. study where they used SMART-seq on a retinal bipolar cell enriched population of cells^21^. Even though our ML algorithm was trained only on droplet-based data, our algorithm labels most of this dataset as retinal bipolar cells, with the next most common cell types being amacrine. The same result was found by Shekhkar et al. (Supplemental Figure 5). To more generally evaluate performance we use the precision recall (PR) curve, which visualizes the ability of the model to precisely label known cells at a given confidence. The area under the PR curve (AUC) summarizes the effectiveness of the model across different cell types, with 1 being the highest performance. The xgboost model can predict rods, bipolar cells, and Müller glia with near perfect performance (Supplemental Figure 6). Most of the remaining cell types can be predicted with an AUC over 0.9 Supplemental Table 2. Several of the precursor cell types (photoreceptor, AC/HC, and neurogenic) were labeled with lower confidence Supplemental Table 3. The next most common labels for these cell types were either other precursor cells (e.g. 100 AC/HC Precursors were labeled as Neurogenic) or the adjacent terminal cell type (e.g. 254 photoreceptor precursors were labeled as cones). Other cell types that were challenging for the model to predict were the artery, choriocapillaris, and vein. Artery, choriocapillaris, and vein are constructed from endothelial cells and we find that for all three of these, endothelial was the second most common label. Overall, our xgboost based ML model shows strong accuracy across the major cell types of the retina (overall AUC of 0.98).

### ML cell type labels result in high study diversity for each cell type

After ML projection of cell type labels from the original 375966 labels onto a total of 758278 cells we hav substantially improved the number of studies per cell type. For example, we went from 8 human studies with labele Müller Glia to 14 after labeling (Supplemental Table 4). Overall we go from an average of 4 studies per human cell type to 8 and 2 studies per mouse cell type to 8 after transferring the cell type labels.

As another check on the quality of the cell type assignments, we ran the cell and cluster independent haystack gene search and pairwise differential expression tests between the predicted cell types (see methods for further details).^47^ We show the five most differentially expressed genes for each of the major retina cell types are consistent with known retinal cell markers (Figure 3a). As a simple metric to identify known and unknown genes relating to the cell type specific expression we search PubMed for the number of publications with two searches per gene. We expect most of the genes identified to be known in the literature. The first search is the more precise “gene AND cell type” (e.g. “PDE6H AND Cones”) and the second search is the more inclusive “gene AND retina” (e.g. “PDE6H AND Retina”). Of the 50 genes in (Figure 3a), 37 had one or more citations in the gene - cell type search (Supplemental File celltype_markers.tsv) and 45 had one or more citation in the inclusive search. The 50 genes had a mean of 46 studies (with the inclusive gene by “retina” search). In contrast, 100 randomly chosen genes had a mean of 2 (wilcox test p < 1.44 x 10**-17).

### The scVI-based scEiaD UMAP projection blends batches and species while separating cell types

The 2D UMAP projection of the scVI-calculated batch corrected 8 latent dimensional space blends the 33 studies together while also maintaining distinct space for the 31 unique cell types. We also see good mixing across all the droplet and well based single cell technologies (Supplemental Figure 7). We see the neurogenic and progenitor populations from which the retinal cell types are derived near the center of the UMAP visualization. The photoreceptor precursors are adjacent to the neurogenic population and, as demonstrated by the RNA velocity dynamics, flow into the rods and cones. The amacrine and horizontal precursors (AC/HC) likewise flow from the neurogenic center into the mature amacrine and horizontal cells.

The photoreceptors (cones and rods) of the retina which are responsible for color and low-light vision, respectively, are near each other in the UMAP space. The major remaining retina cell types, by proportion in the mammalian eye are the Müller Glia, which are a glial cell type which help support the neurons of the retina. Next we have the neural cell types which transmit and help interpret the signals from the photoreceptors before they leave the retina via the optic stalk: the amacrine cells, retinal ganglia, horizontal, and bipolar cells. All of these cells are in well separated spaces in the UMAP. Finally we see across species that the major cell types overlap each other (Figure 3d). The macaque retina cells are not present in the precursor/neurogenic center of the UMAP as expected because only fully developed tissues were sampled in these data.

While by eye the UMAP 2D projection generally blends together the three different species (macaque, mouse, human) in the UMAP visualization, we more rigorously tested this by changing the inputs to our xgboost cell type predictor machine learning system. We trained two new models: one with only human data and one with only mouse data. We then applied this model to the other two species to see whether the model trained on one species was generally transferable to another species. Again, both models (human only and mouse only) both proved to be highly accurate at predicting cell types in the other species with the terminal cell types (Supplemental Figure 8.

### Projection of outside data onto scEiaD demonstrates similarities and differences of iPSC and organoids to primary cells

The hard work of creating this resource can be leveraged and extended by the wider community with a few relatively simple steps and modest compute requirements. Very briefly, if outside groups quantify their mouse or human scRNA with kallisto (bustools) and the same references (see methods), they can overlay their data on top of scEiaD by 1. installing scVI (version 0.9.0 or higher), 2. downloading our 13 megabyte scVI model, and 3. following the Jupyter notebook on Google colab that we provide as a live demo available from https://github.com/davemcg/scEiaD. We demonstrate the power of this approach in two ways.

First, a wild-type retinal organoid dataset from Kallman et al was projected onto scEiaD.^48^ In the Google colab notebook we show how a subset of the reads from Kallman et al.’s SRR12130660 can be processed from the raw reads to a UMAP visualization in under 10 minutes. Indeed we see how organoid-based retinal cell types can be detected overlapping primary cells for RPCs, photoreceptors, amacrine cells, and retinal ganglion cells (Supplemental Figure 9).

Finally, we demonstrate how the RPE derived from the Bharti iPSC differentiation process express canonical RPE markers of TTR and RPE65 but end up in a slightly different position in the UMAP projectionFigure 4a). Differential expression analysis (see methods) identified that vimentin is nearly exclusively expressed in the iPSC-based RPE Figure 4b). This result is not surprising as Hunt et al. demonstrated that cultured and proliferating RPE express high levels of vimentin^49^ and this same trend is seen in bulk RNA-seq from stem cell RPE and primary RPE Supplemental Figure 10).^50^

**Figure 4:**
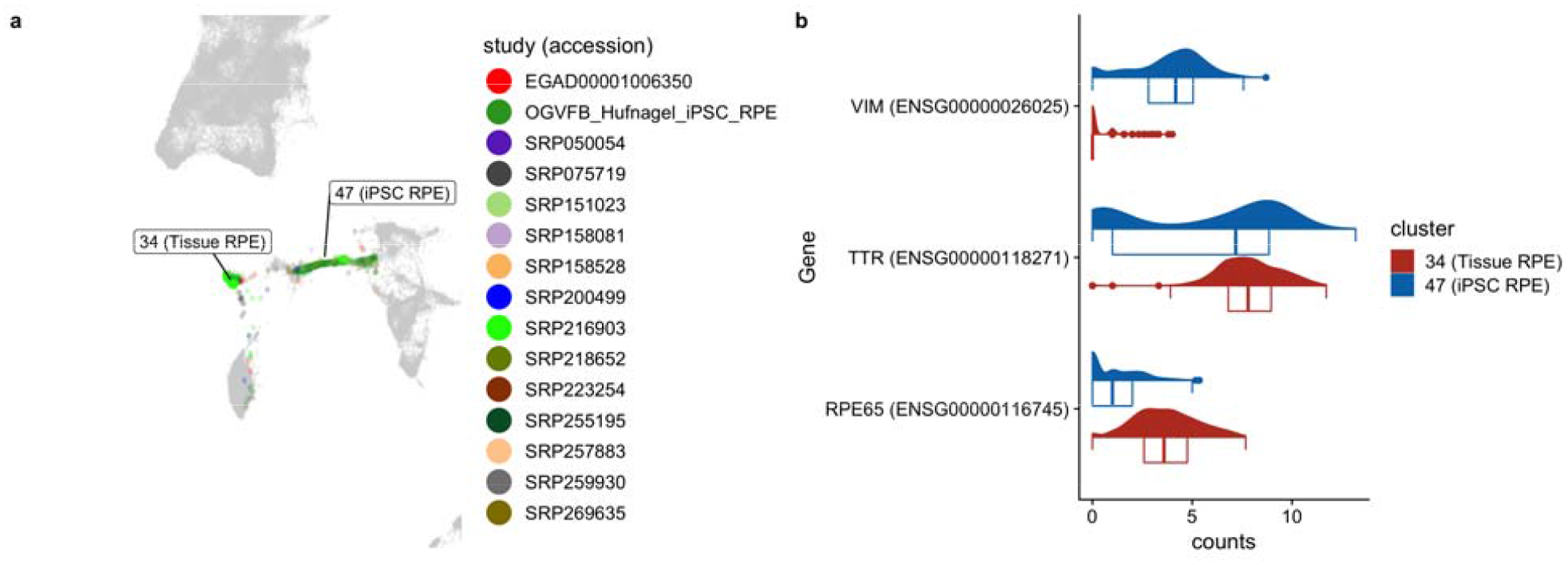
a. RPE distribution colored by study demonstrates how the most RPE are in two locations. iPSC-based RPE we provided are located more enriched in cluster 47. Tissue RPE more enriched in cluster 34 b. Violin plot of two functional RPE markers (TTR, RPE65) and vimentin *(proliferating RPE marker)*

## Methods

### Reproduciblity and data availability

The set of Snakemake pipelines that takes in the raw fastq sequence and outputs the scEiaD is at https://github.com/davemcg/scEiad.51 The publication commit is #ffdf738. Furthermore, the repository has bee deposited at Zenodo under 10.5281/zenodo.5129265 We will briefly discuss the pipeline choices, programs an algorithms, and versions below. For the R packages, we provide package versions as the supplementary file “R_session_info.txt”

We have deposited the 10x (v2) fastq files for our iPSC-based RPE data in GEO at accession GSE180662. We give download links for the count matrices, seurat and anndata objects, cell level metadata, and our codebases i Supplemental Table 5.

### Quantification of gene counts

Gene quantification is handled by the SnakeQUANT snakefile. First, we generated multiple quantification indices to facilitate calculation of RNA velocity. For each droplet technology, we generated a separate set of transcript sequences that contain both exonic and intronic sequences using the “get_velocity_files” function from the R package BUSpaRse. The reference transcriptome annotation used to build sets of transcript sequences were the Gencode “gencode.vM25.annotation.gtf.gz” and “gencode.v35.annotation.gtf.gz” for mouse and human, respectively.^52, 53^ Because the Macaca Fascicularis genome is less well annotated we used the Ensembl release 101 genome and transcriptome annotation “Macaca_mulatta.Mmul_101.gtf.gz.”^54^ A single set of exonic transcript sequences was created for each species for all well techonologies. A kallisto quantification index was generated for each of these fastqs using kallisto index (0.46.2).^55^ Well based samples were quantified using kallisto quant. For the relatively few single ended samples, the params “–single −l 200 -s 30” were used for kallisto quant. Otherwise the “– bias” flag was added. For the droplet-based samples, we adapted the bustools workflow for generating spliced and unspliced count matrices. (0.39.4).^56^

### Intersection of gene names between mouse, macaque, and human

To facilitate comparison of gene expression across species, where possible we converted mouse and macaque gene ids and names to human ones. We downloaded a mapping of orthologous genes between human, mouse, and macaque using the Ensembl BioMart web browser in November 2020. We identified 15,759 human genes that could be directly mapped to mouse and macaque orthologs. Genes present in mouse or macaque that were not found in human were not used for HVG gene selection, but were retained and used for differential gene expression.

### Custom macaque reference quantification

As we noticed that several retina marker genes (e.g. NRL and CRX) had very low expression in the macaque data we quantified the scRNA data twice: once with the Ensembl reference and again with the same Gencode human reference used for the human data. We compared the gene-level counts for each cell and replaced the macaque gene count with the human counts if the human counts were greater than the macaque counts and, to prevent genes with very few total counts from being used, we required the counts greater than the first quartile of non-zero macaque gene expression.

### Remove empty droplets and further QC

After bustools count, we used R (3.6.2) to remove empty droplets. The BUSpaRse package was used to input the bustools counts mtx file. The DropletUtils package with the “barcodeRanks” function was used to automatically detect the inflection point in the barcode count ranks that delineates the likely empty droplets.^57^ We then removed cells with percent mitochondrial reads of >10%. After merging the individual count matrices into one sparse matrix, we created a Seurat version 3 object and removed cells with fewer than 200 detected unique genes, and for the droplet data, more than 3000 detected genes (these are more likely to be doublets).^32^

### Normalization and batch effect correction

The following steps (normalization through benchmarking are handled by the SnakePOP pipeline) We tested several gene count normalization approaches as we were not certain which would produce an optimal outcome: standard (default Seurat, library size normalization, then log transform), sqrt (same, but with sqrt normalization), libSize (omit the log or sqrt normalization), scran, SCT from Seurat, and for scVI, no normalization (counts).^58^ Our R implementation of the normalization approaches as well as how we constructed the Seurat v3 object can be found in the supplementary file “make_seurat_obj_functions.R”

### Batch normalization under a grid search procedure

We tested scArches, bbknn, insct, magic, scVI, CCA, scanorama, harmony, fastMNN, combat, none against 2000 HVGs, the different gene count normalization procedures discussed above, and both 8 and 30 outputted batch corrected latent dimensions. The latent dimensions are the input for clustering, the 2D UMAP visualization, and the xgboost machine learning to transfer cell type labels to unlabeled cells. We were unable to run every method successfully with every normalization method. For example, magic could not complete with the standard or libSize normalization. Insct was only able to complete the scran normalization. We also tried the DESC, liger, and Conos batch corrections methods but were unable to get them to work reliably so they were dropped. While we attempted to use “default” parameters wherever possible, we had to deviate from this to get Seurat’s CCA procedure to complete. CCA reciprocally tries to integrate all batches and even with a subset of cells we regularly got “long vector” errors. We got around this issue by setting the human batches as the “reference.” The batch correction steps implementation can be found the supplementary file “merge_methods.R” and in the github repo (https://github.com/davemcg/scEiaD/blob/master/src/merge_methods.R).

### Clustering and UMAP

Louvain-Jaccard clustering against the batch corrected latent dimensions used the Seurat implementation.^59^ For the all methods benchmarking we used k-nearest neighbors (knn) of 20. For the scVI-only tuning reduced the knn parameters to 5 and 7 (where 5 gives more clusters than 7) to increase the cluster number. We also used the leiden algorithm as implemented by PARC with a resolution of 0.6 and 0.8 (higher results in more clusters).^60, 61^ These two resolutions were chosen as they roughly gave the same number of clusters at the Seurat Louvain-Jaccard approach with a knn 7.

The UMAP visualization was calculated with the Seurat “RunUMAP” using the uwot R package.^62^ We tried min.dist parameters of 0.001, 0.1, and 0.3 and tried n.neighbors across 15, 30, 100, and 500. A smaller min.dist value gives “tighter” groupings while a higher number of n.neighbors uses a larger number of near cells to calculate the global positioning.

### Benchmarking and scPOP

We wrote the scPOP R package to unify the LISI and Silhouette metrics from Harmony and kBet, respectively, along with NMI and ARI.^36, 63^ LISI and Silhouette require a dense matrix, which is a problem for our data as a 766615 cell by 8 latent dimension dense matrix cannot fit in out largest available compute node (1.5 TB). We down-sampled the dataset to ∼100,000 cells, taking care to keep all rarer cell types for the LISI and Silhouette benchmarking.

To merge these metrics into a balanced single score, we Z scale each and sum them. scPOP produces both tables and visualizations allowing the user to quickly see both the interplay of batch mixing and cluster/cell type separation and the overall performance. If a user wishes to prioritize batch mixing or cluster/cell type separation we let the user provide a custom batch/cluster-cell type scaling value (1 is the default).

### Multi-step doublet removal

To identify probable doublets (more than one cell in a droplet) we ran DoubletDetect and scrublet and calculated the distribution of DoubletDetect and scrublet scores across all clusters and removed clusters with a score in both metrics greater than 4 standard deviations above the mean.^64, 65^ This removed another 23,457 cells, leaving 766,615 in total.

### xgboost based cell type model

In order to identify cell types for the 361,456 unlabeled cells we designed a custom xgboost based cell type classifier. We took labeled data and split it into training (2/3) and test (1/3) sets, stratified by cell type. The input features used to train the model are the scVI latent dimensions, the total number of reads in each cell, the number genes detected in each cell, and the percent mitochondrial gene expression of each cell. We additionally generated features using the age of each sample by group sample into three developmental categories (Early Development, Late Development, and Adult) and then generated a one-hot encoded feature for each category. In order to speed up training times, we used the gpu implementation of the xgboost algorithm from the the xgboost python library. The model was trained using default parameters. The trained model had an overall macro and micro AUC score of 0.98 and 0.99, respectively. This model was then used to identify labels for all cells. Unlabeled data was pre-processed identically to training data and fed into model to generate a vector of label probabilities for each cell. We selected the highest label probability for each cell, and required a minimum probability of 0.5 to assign a label to a cell.

For the organism specific xgboost ML we followed the above procedure, except that we combined Early RPCs, Late RPCs, and RPCs into one category and did not attempt to predict the non-retina cell types (e.g. fibroblasts) as there were very few labeled cells across all three organisms.

### Marker gene identification

To identify marker genes across the CellType (predict) and cluster groups, we used the scran findmarkers (wilcox test) along with the singleCellHaystack algorithm.^47^ The scran findmarkers test runs a wilcox test in a pairwise manner (e.g. Rods vs all other cell types). It returns an overall p-value (and FDR) that assesses how well the gene is at separating the group of interest from all other cells. It also returns for each pair-wise comparison an area under the curve (AUC) score, where 1 is a perfect power to distinguish and 0 is no power. The singleCellHaystack algorithm uses a Kullback-Leibler divergence (D KL) measurement of the scVI lower dimensional space to identify genes with non-random distribution. A higher D KL score represents a gene with “specific” expression in the lower dimensional space and is used to calculate a FDR corrected p value against the full distribution of D KL values. We filtered to keep genes with scran FDR < 1, a mean AUC > 0.2, and a log10(D KL FDR) < −10000. No more than 50 genes for each cell type were retained (sorted by mean AUC).

### Calculation of RNA velocity

RNA velocity calculations were with the velocriaptor wrapping of the scVelo python library.^66^ From the anndata objects generated by our Snakemake pipeline we calculated velocity across all genes. Genes without detectable velocity were dropped. The scVI generated latent space (instead of PCA) was used to calculate first and second order moments. The calculated moments were used to estimate RNA velocity. Differential velocity was tested between celltypes using pairwise wilcox rank sum tests.

## Conclusion

### Limitations

The scVI model is first built the on human data. The mouse and macaque data are then projected (or queried) onto it with the scVI implementation of the scArches method. While this system works very well to integrate information between these three species, this approach may not scale to more distantly related species. Another limitation is that discrepancies between cell type labels between different labs makes certain transitioning cell type labels a bit imprecise. One example is how the rods and photoreceptor precursor labels partially overlap. Though we attempted to ameliorate the issue by removing cell type labels in large disagreement with the consensus, some disagreements could propagate into our machine labeled cells type assignments. These issues may reflect labeling continuous processes with discrete labels.

While scEiaD distinguishes the major cell types very well, some of the cell types contain many “sub types” - notably the amacrine cells have a huge variety in morphology with a single cell based study identifying over sixty different types of amacrine cells in mouse.^29^ At this time our batch corrected pan retina cell space does not precisely resolve these sub cell types with high resolution. We are actively working to “sub cluster” the cell types so we can robustly and reliably identify the high diversity of retinal cell types across the entire retina.

### The scEiaD is a unique ocular resource that provides a highly diverse, large N dataset with a relatively small amount of compute power

We have assembled the largest ocular single cell transcriptome database to date. The rapid of advancement of algorithms to batch correct and process data continue to reduce the computational requirements to handle huge numbers of cells. The scVI batch correction step with around one million cells runs within an hour on a GPU and 150GB of memory. This places this crucial step within the capabilities of a moderately powerful computer or a cloud compute node. Further downstream processing can largely be done a computers with 64+ GB of memory and a few hundred GB of disk space. We believe that our efforts can be replicated in any other tissue / system with a large number of independent studies by a small number of computational scientists following our general approach. We provide the completed analysis as both Scanpy (h5ad) and Seurat objects (Supplemental Table 5).

### Benchmarking and quantitation of integration performance is crucial for meta-atlas studies

We originally intended to use the Seurat CCA method to integrate the datasets. However, the long run-times of CCA and poor integration of our known cell types led us to benchmark more methods and parameters. After adding more integration methods we first attempted “hand-assess” the integration results by using the UMAP 2D projection view. This proved to scale poorly and this led us to curate some of the more useful benchmarking algorithms (NMI, ARI, silhouette, and LISI) that roughly matched our “hand-assessed” results into the scPOP R package. While we chose the scVI algorithm, we strongly suggest any other groups attempting a similar meta-atlas construction chose a quantifiable set of criteria so optimal methods and parameters can be picked. We found in our analysis that substantial differences in the integration performance can come from altering the number of HVGs and the number of latent dimensions.

### Transfer of cell type labels from a smaller number of studies onto the remainining cells is a powerful way to increase diversity in a meta-atlas

We first hand curated over 350,000 published cell type labels across 24 publications. With a xgboost algorithm using the latent dimensions, cell age classification (developing or matured), and the UMAP coordinates, we can very accurately label the remaining cells. Many other cell type labeling algorithms and systems exist for those groups less willing or able to tune a machine learning algorithm. For example, the developers of scVI also have a cell type label projection algorithm called scANVI.^67^ Whichever approach you use, taking a smaller number of high quality labels and projecting them onto the remaining cells is a powerful way to leverage community knowledge across a huge diverse dataset.

### Projection allows community knowledge to be leveraged by all

Many retina atlases have been published to date. We argue that we have created the first atlas that is generally useful because 1. our dataset/atlas is several times larger than any other published set, 2. our data is available via download in several forms at https://github.com/davemcg/scEiaD and Zenodo accession 5129265, and crucially 3. we provide a Google colab/Jupyter notebook which step-by-step demonstrates out how to use scVI to project (or query) outside data onto our scEiaD with minimum compute resources. We demonstrate concretely how this can work by showing how iPSC-based RPE can be queried onto the reference dataset to demonstrate both similarities and dissimilarities in their transcriptomes.

## Supporting information

Celltype markers

kallisto pseudoalignment stats

R 3.6 session info

R 4.0 session info

UMAP all methods, colored by study accession

UMAP all methods, colored by cell type

UMAP all methods, colored by organism

Seurat obj creation and integration implementation scripts

## Supplemental Information

**Supplemental Table 1:**
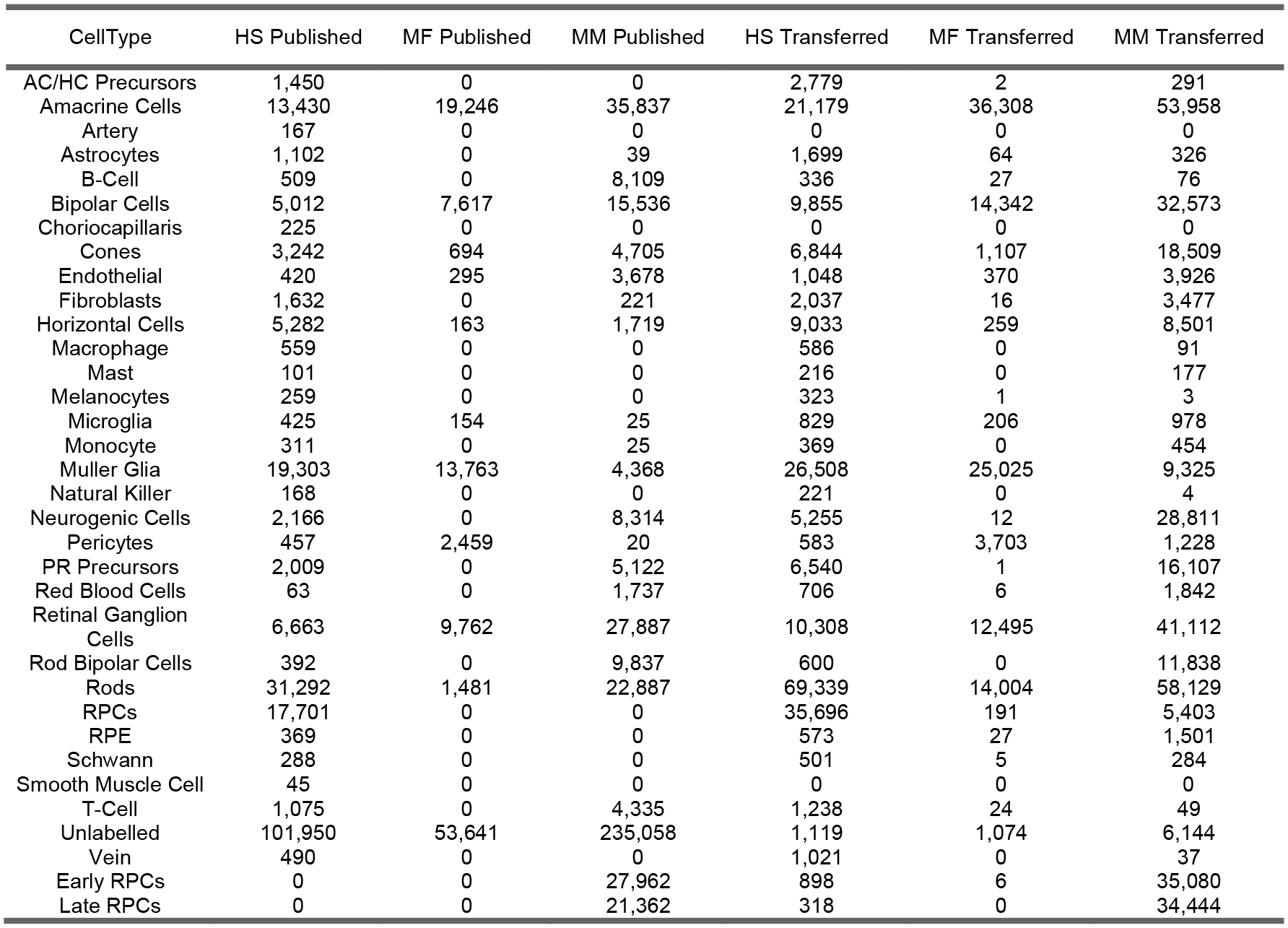
Counts for cell type labels. Published are the author created labels from the published datasets. Transferred are the cell labels that were transferred by our xgboost-based machine learning model onto the entire scEiaD dataset.

**Supplemental Table 2:** Area under the precision recall curve (AUC) for each cell type, split by study. The “All” study is the AUC score across all cells within the cell type

| AUC | Cell Type | Study |
| --- | --- | --- |
| 0.76 | AC/HC_Precurs | All |
| 0.71 | AC/HC_Precurs | SRP151023 |
| 0.83 | AC/HC_Precurs | SRP223254 |
| 0.99 | Amacrine Cells | All |
| 0.20 | Amacrine Cells | E-MTAB-7316 |
| 1.00 | Amacrine Cells | EGAD00001006350 |
| 0.97 | Amacrine Cells | SRP050054 |
| 0.98 | Amacrine Cells | SRP075719 |
| 0.98 | Amacrine Cells | SRP151023 |
| 0.93 | Amacrine Cells | SRP158081 |
| 1.00 | Amacrine Cells | SRP158528 |
| 1.00 | Amacrine Cells | SRP194595 |
| 1.00 | Amacrine Cells | SRP200499 |
| 0.72 | Amacrine Cells | SRP222001 |
| 1.00 | Amacrine Cells | SRP222958 |
| 0.98 | Amacrine Cells | SRP223254 |
| 1.00 | Amacrine Cells | SRP255195 |
| 0.97 | Astrocytes | All |
| 1.00 | Astrocytes | E-MTAB-7316 |
| 0.97 | Astrocytes | EGAD00001006350 |
| 0.94 | Astrocytes | SRP050054 |
| 0.94 | Astrocytes | SRP200499 |
| 1.00 | Astrocytes | SRP255195 |
| 0.97 | B Cell | All |
| 0.81 | B Cell | EGAD00001006350 |
| 1.00 | B Cell | SRP131661 |
| 0.41 | B Cell | SRP218652 |
| 0.91 | B Cell | SRP257883 |
| 0.98 | Bipolar Cells | All |
| 1.00 | Bipolar Cells | E-MTAB-7316 |
| 1.00 | Bipolar Cells | EGAD00001006350 |
| 0.97 | Bipolar Cells | SRP050054 |
| 0.99 | Bipolar Cells | SRP075719 |
| 0.97 | Bipolar Cells | SRP151023 |
| 0.94 | Bipolar Cells | SRP158081 |
| 1.00 | Bipolar Cells | SRP158528 |
| 1.00 | Bipolar Cells | SRP194595 |
| 1.00 | Bipolar Cells | SRP222001 |
| 1.00 | Bipolar Cells | SRP222958 |
| 0.98 | Bipolar Cells | SRP223254 |
| 1.00 | Bipolar Cells | SRP255195 |
| 0.94 | Cones | All |
| 1.00 | Cones | E-MTAB-7316 |
| 1.00 | Cones | EGAD00001006350 |
| 0.96 | Cones | SRP050054 |
| 0.71 | Cones | SRP151023 |
| 0.88 | Cones | SRP158081 |
| 0.97 | Cones | SRP158528 |
| 1.00 | Cones | SRP194595 |
| 1.00 | Cones | SRP200499 |
| 1.00 | Cones | SRP222001 |
| 0.47 | Cones | SRP222958 |
| 0.97 | Cones | SRP223254 |
| 0.90 | Cones | SRP255195 |
| 0.98 | Early RPCs | All |
| 0.99 | Early RPCs | SRP158081 |
| 0.97 | Endothelial | All |
| 0.98 | Endothelial | EGAD00001006350 |
| 0.99 | Endothelial | SRP050054 |
| 1.00 | Endothelial | SRP131661 |
| 0.94 | Endothelial | SRP158528 |
| 1.00 | Endothelial | SRP194595 |
| 0.96 | Endothelial | SRP200499 |
| 0.00 | Endothelial | SRP218652 |
| 1.00 | Endothelial | SRP222958 |
| 0.59 | Endothelial | SRP255195 |
| 0.97 | Fibroblasts | All |
| 1.00 | Fibroblasts | EGAD00001006350 |
| 0.95 | Fibroblasts | SRP050054 |
| 0.99 | Fibroblasts | SRP131661 |
| 0.53 | Fibroblasts | SRP218652 |
| 0.99 | Fibroblasts | SRP257883 |
| 0.96 | Horizontal Cells | All |
| 1.00 | Horizontal Cells | E-MTAB-7316 |
| 1.00 | Horizontal Cells | EGAD00001006350 |
| 0.97 | Horizontal Cells | SRP050054 |
| 0.99 | Horizontal Cells | SRP151023 |
| 0.79 | Horizontal Cells | SRP158081 |
| 0.92 | Horizontal Cells | SRP158528 |
| 0.95 | Horizontal Cells | SRP200499 |
| 1.00 | Horizontal Cells | SRP222001 |
| 1.00 | Horizontal Cells | SRP222958 |
| 0.98 | Horizontal Cells | SRP223254 |
| 1.00 | Horizontal Cells | SRP255195 |
| 0.97 | Late RPCs | All |
| 0.97 | Late RPCs | SRP158081 |
| 0.70 | Macrophage | All |
| 0.58 | Macrophage | SRP218652 |
| 0.97 | Macrophage | SRP257883 |
| 0.83 | Mast | All |
| 0.92 | Mast | EGAD00001006350 |
| 0.96 | Mast | SRP218652 |
| 0.97 | Melanocytes | All |
| 1.00 | Melanocytes | EGAD00001006350 |
| 0.00 | Melanocytes | SRP218652 |
| 0.98 | Melanocytes | SRP257883 |
| 0.96 | Microglia | All |
| 0.95 | Microglia | E-MTAB-7316 |
| 1.00 | Microglia | EGAD00001006350 |
| 0.96 | Microglia | SRP050054 |
| 0.96 | Microglia | SRP158528 |
| 1.00 | Microglia | SRP194595 |
| 0.79 | Microglia | SRP200499 |
| 1.00 | Microglia | SRP222001 |
| 0.90 | Microglia | SRP222958 |
| 1.00 | Microglia | SRP255195 |
| 0.97 | Monocyte | All |
| 0.99 | Monocyte | EGAD00001006350 |
| 0.99 | Monocyte | SRP200499 |
| 0.99 | Muller Glia | All |
| 1.00 | Muller Glia | E-MTAB-7316 |
| 1.00 | Muller Glia | EGAD00001006350 |
| 1.00 | Muller Glia | SRP050054 |
| 1.00 | Muller Glia | SRP075719 |
| 0.80 | Muller Glia | SRP151023 |
| 0.74 | Muller Glia | SRP158081 |
| 1.00 | Muller Glia | SRP158528 |
| 1.00 | Muller Glia | SRP194595 |
| 1.00 | Muller Glia | SRP200499 |
| 1.00 | Muller Glia | SRP222001 |
| 0.99 | Muller Glia | SRP222958 |
| 0.99 | Muller Glia | SRP223254 |
| 1.00 | Muller Glia | SRP255195 |
| 0.89 | Natural Killer | All |
| 0.97 | Natural Killer | EGAD00001006350 |
| 0.81 | Neurogenic Cells | All |
| 0.76 | Neurogenic Cells | SRP151023 |
| 0.85 | Neurogenic Cells | SRP158081 |
| 0.63 | Neurogenic Cells | SRP223254 |
| 0.95 | Pericytes | All |
| 0.97 | Pericytes | EGAD00001006350 |
| 0.98 | Pericytes | SRP158528 |
| 0.14 | Pericytes | SRP218652 |
| 0.98 | Pericytes | SRP257883 |
| 0.69 | Photoreceptor Precursors | All |
| 0.48 | Photoreceptor Precursors | SRP151023 |
| 0.91 | Photoreceptor Precursors | SRP158081 |
| 0.40 | Photoreceptor Precursors | SRP223254 |
| 0.98 | Red Blood Cells | All |
| 1.00 | Red Blood Cells | SRP131661 |
| 0.96 | Red Blood Cells | SRP158081 |
| 0.78 | Red Blood Cells | SRP257883 |
| 0.99 | Retinal Ganglion Cells | All |
| 0.10 | Retinal Ganglion Cells | E-MTAB-7316 |
| 1.00 | Retinal Ganglion Cells | EGAD00001006350 |
| 1.00 | Retinal Ganglion Cells | SRP050054 |
| 0.98 | Retinal Ganglion Cells | SRP151023 |
| 0.93 | Retinal Ganglion Cells | SRP158081 |
| 1.00 | Retinal Ganglion Cells | SRP158528 |
| 0.94 | Retinal Ganglion Cells | SRP200499 |
| 1.00 | Retinal Ganglion Cells | SRP222958 |
| 0.98 | Retinal Ganglion Cells | SRP223254 |
| 1.00 | Retinal Ganglion Cells | SRP255195 |
| 0.99 | Rod Bipolar Cells | All |
| 1.00 | Rod Bipolar Cells | EGAD00001006350 |
| 1.00 | Rod Bipolar Cells | SRP075719 |
| 0.99 | Rod Bipolar Cells | SRP200499 |
| 0.99 | Rods | All |
| 1.00 | Rods | E-MTAB-7316 |
| 1.00 | Rods | EGAD00001006350 |
| 0.97 | Rods | SRP050054 |
| 0.96 | Rods | SRP151023 |
| 0.97 | Rods | SRP158081 |
| 0.96 | Rods | SRP158528 |
| 1.00 | Rods | SRP194595 |
| 1.00 | Rods | SRP200499 |
| 1.00 | Rods | SRP222001 |
| 1.00 | Rods | SRP222958 |
| 0.97 | Rods | SRP223254 |
| 0.96 | Rods | SRP255195 |
| 0.98 | RPCs | All |
| 0.99 | RPCs | SRP151023 |
| 0.99 | RPCs | SRP223254 |
| 0.94 | RPE | All |
| 1.00 | RPE | EGAD00001006350 |
| 0.81 | Schwann | All |
| 0.38 | Schwann | SRP218652 |
| 1.00 | Schwann | SRP257883 |
| 0.97 | T Cell | All |
| 0.98 | T Cell | EGAD00001006350 |
| 0.99 | T Cell | SRP131661 |
| 0.05 | T Cell | SRP218652 |
| 0.99 | T Cell | SRP257883 |
| 0.91 | Vein | All |
| 1.00 | Vein | SRP257883 |

**Supplemental Table 3:**
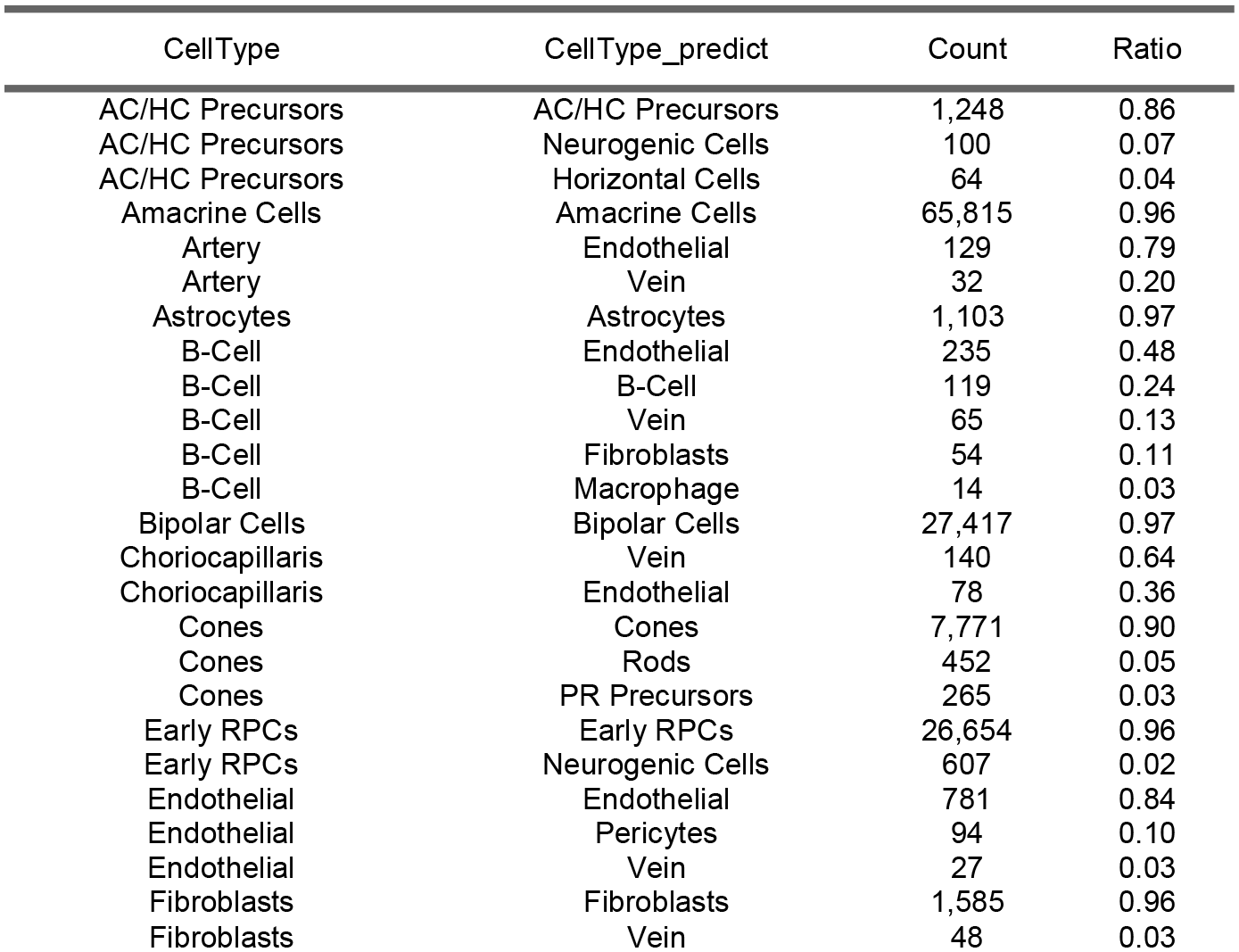
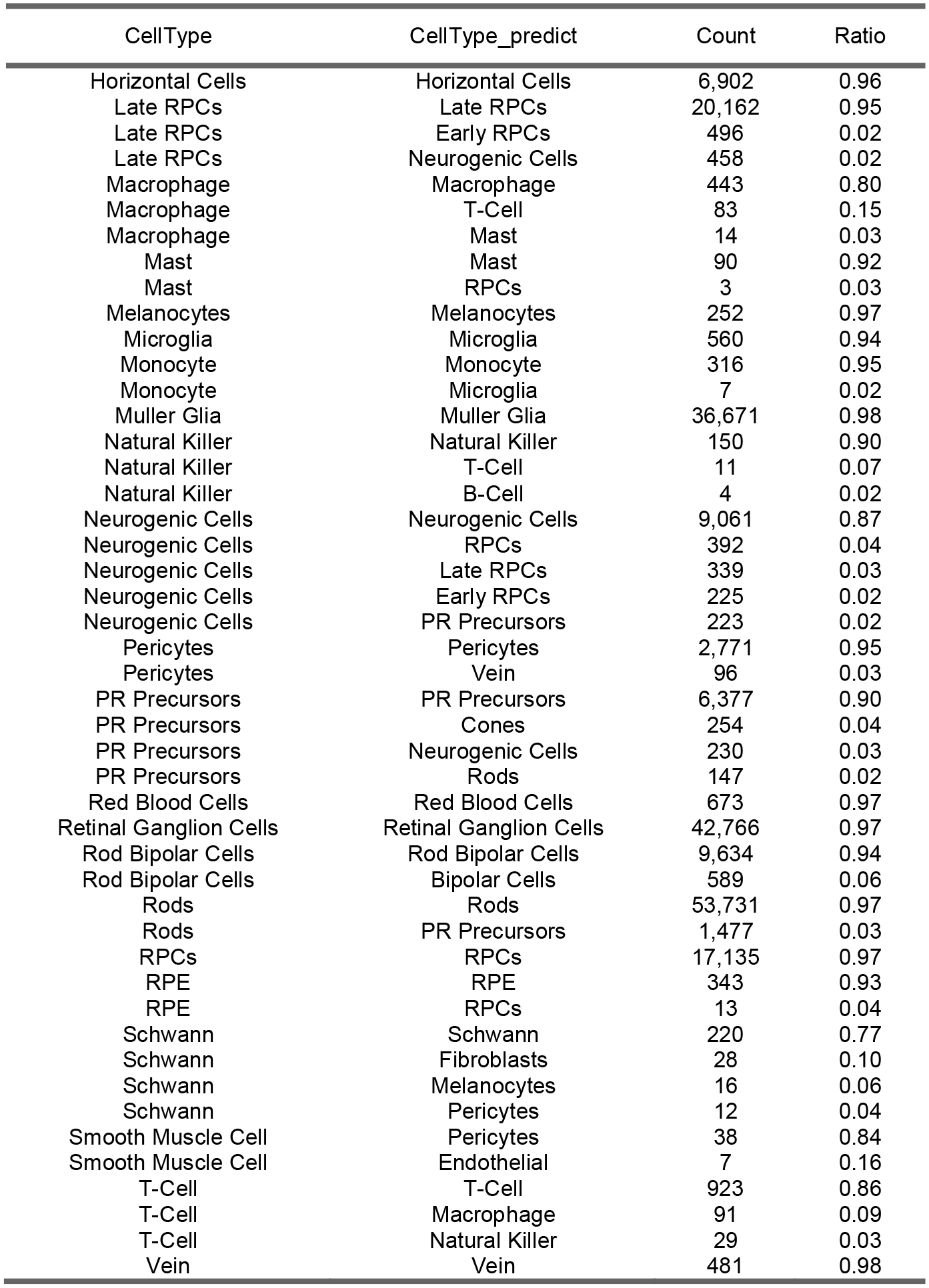
Counts of cell type labels with our xgboost machine learning system (PredCellType) and the published cell type labels (TrueCellType). Ratio is calculated as CellType that were labeled as CellType_predict. Ratio < 0.05 were filtered out from view in the table.

**Supplemental Table 4:**
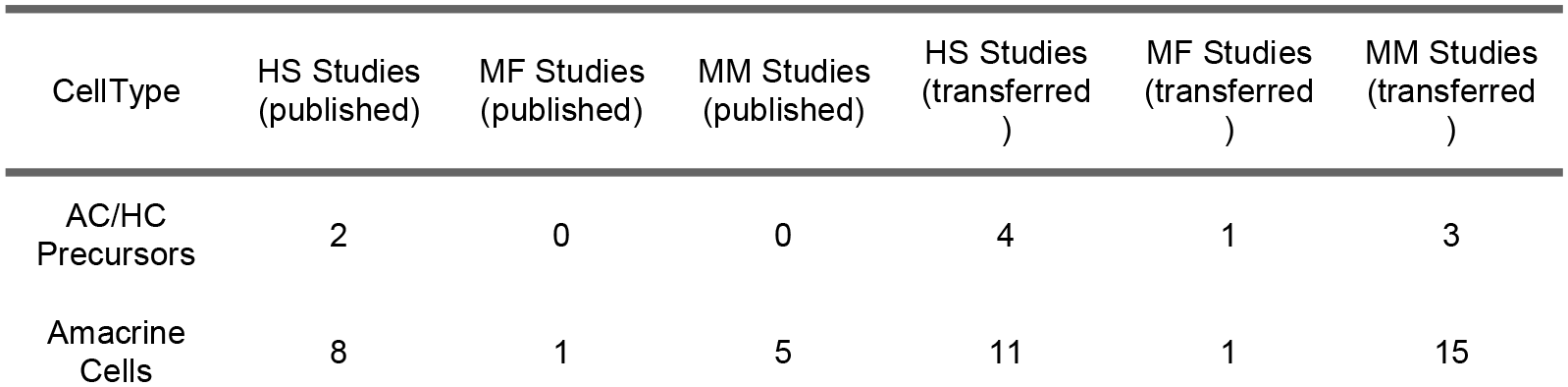
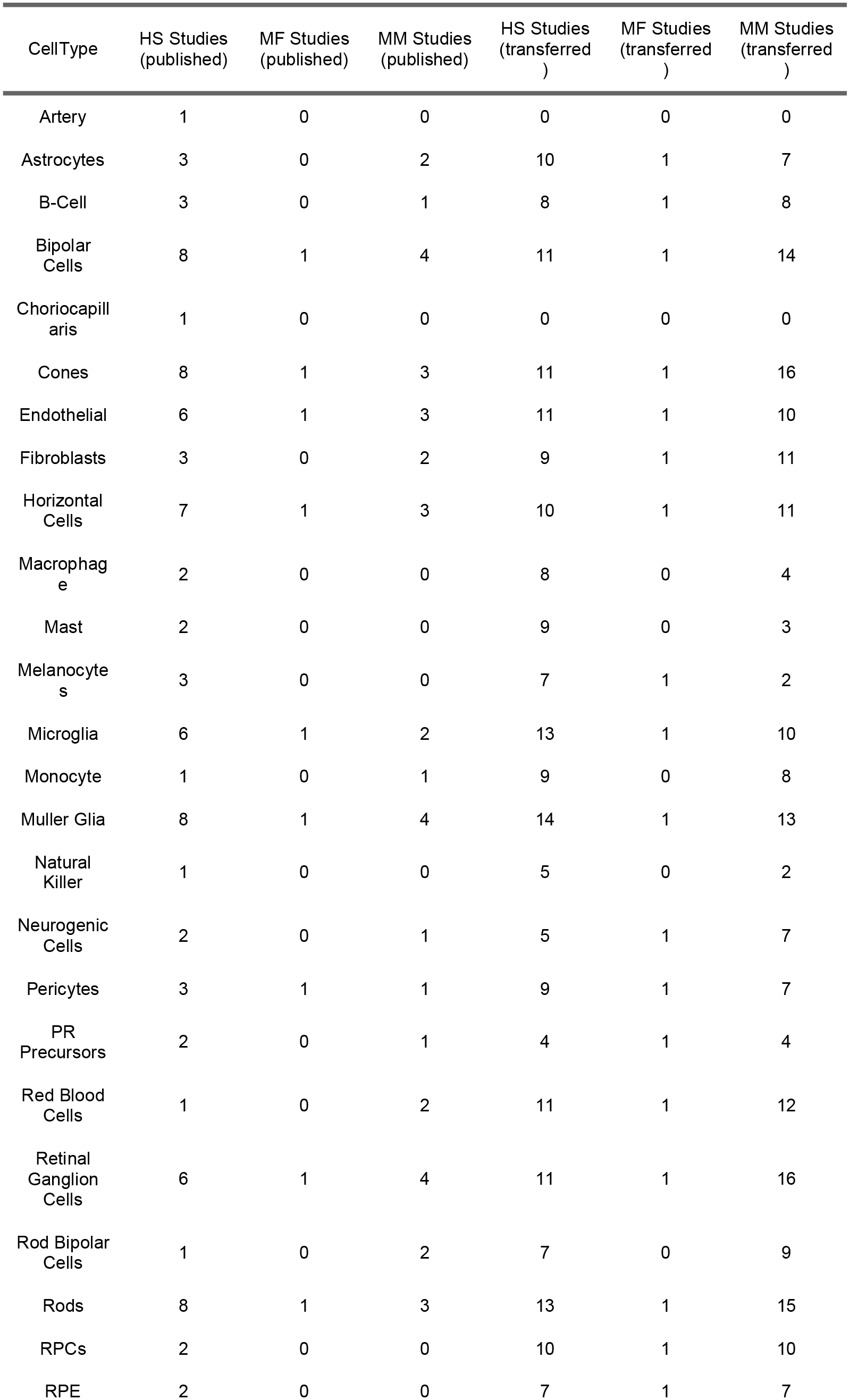
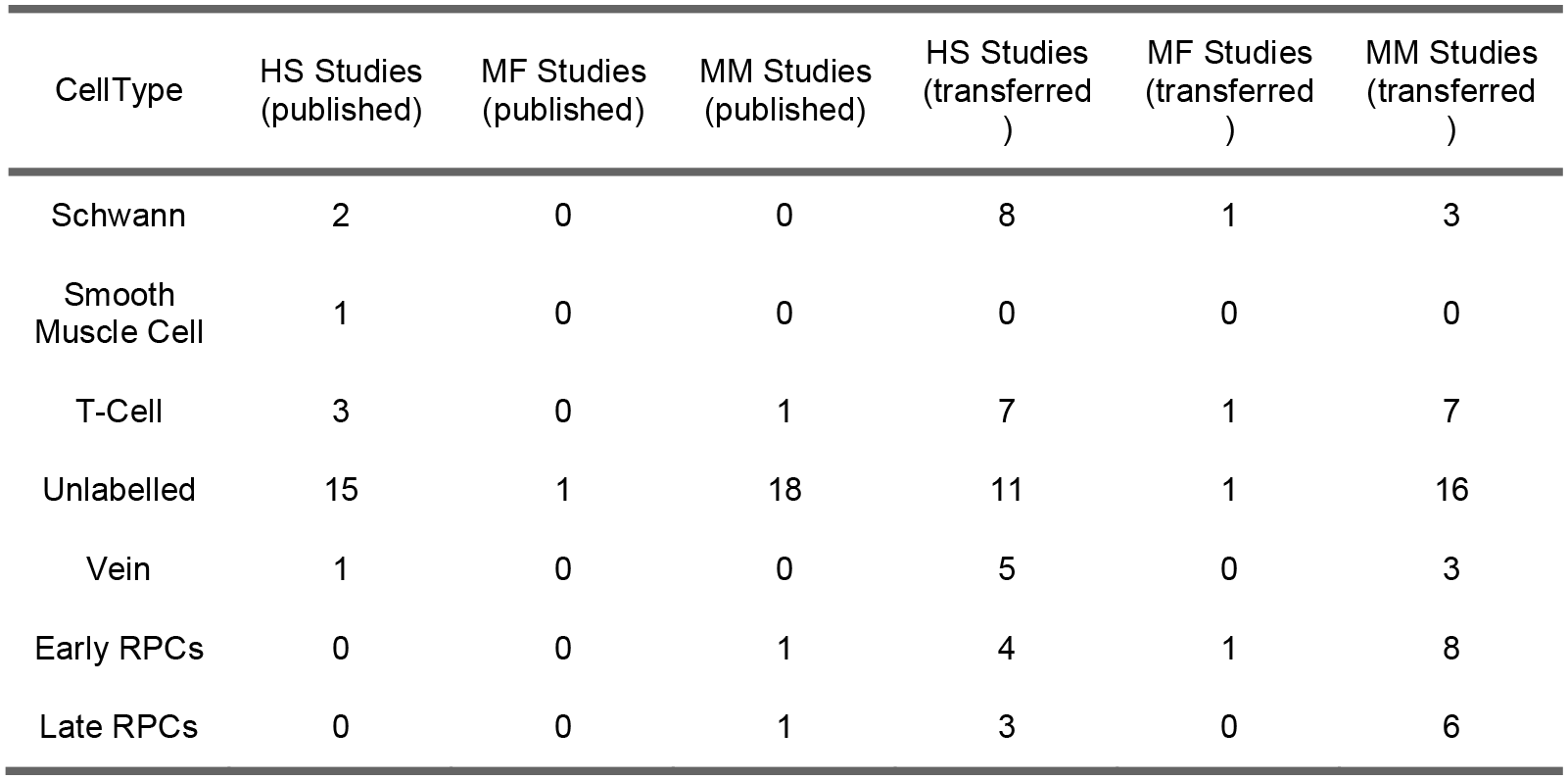
Counts for number of studies with cell types labels before and after cell type label transfer

**Supplemental Table 5:** Links to code and resources

| Resource or File | Link | Accession |
| --- | --- | --- |
| Seurat Object | http://hpc.nih.gov/~mcgaugheyd/scEiaD/2021_03_17/scEiaD_all_seurat_v3.Rdata |  |
| Anndata Object | http://hpc.nih.gov/~mcgaugheyd/scEiaD/2021_03_17/scEiaD_all_anndata.h5ad |  |
| Counts, R sparse matrix | http://hpc.nih.gov/~mcgaugheyd/scEiaD/2021_03_17/counts.Rdata |  |
| Cell level metadata | http://hpc.nih.gov/~mcgaugheyd/scEiaD/2021_03_17/metadata_filter.tsv.gz |  |
| Codebase for scEiaD creation | https://github.com/davemcg/scEiaD | ffdf738 |
| Codebase for scEiaD manuscript | https://github.com/davemcg/scEiaD_manuscript |  |
| Zenodo deposit of all above files and code | https://zenodo.org/record/5129265 | 5129265 |
| iPSC RPE scRNA Raw Fastq Files | https://www.ncbi.nlm.nih.gov/geo/query/acc.cgi?acc=GSE180662 | GSE180662 |

**Supplemental Figure 1:**
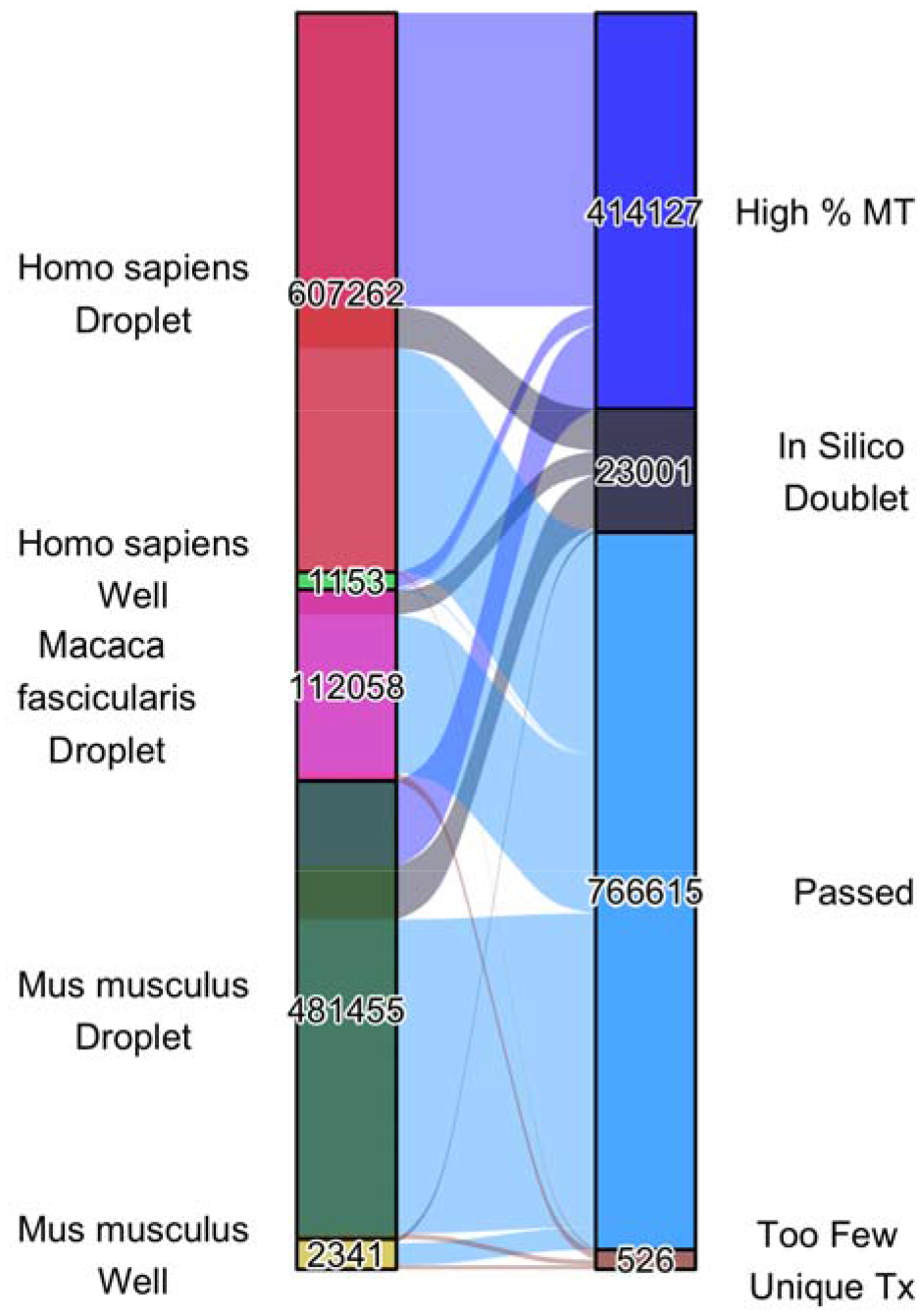
The left bar delineates the number of cells for each organism - technology combination. The right bar specifies the number of each cells in each post QC category. In silico doublets were identified with scrublet and DoubletDetector.

**Supplemental Figure 2:**
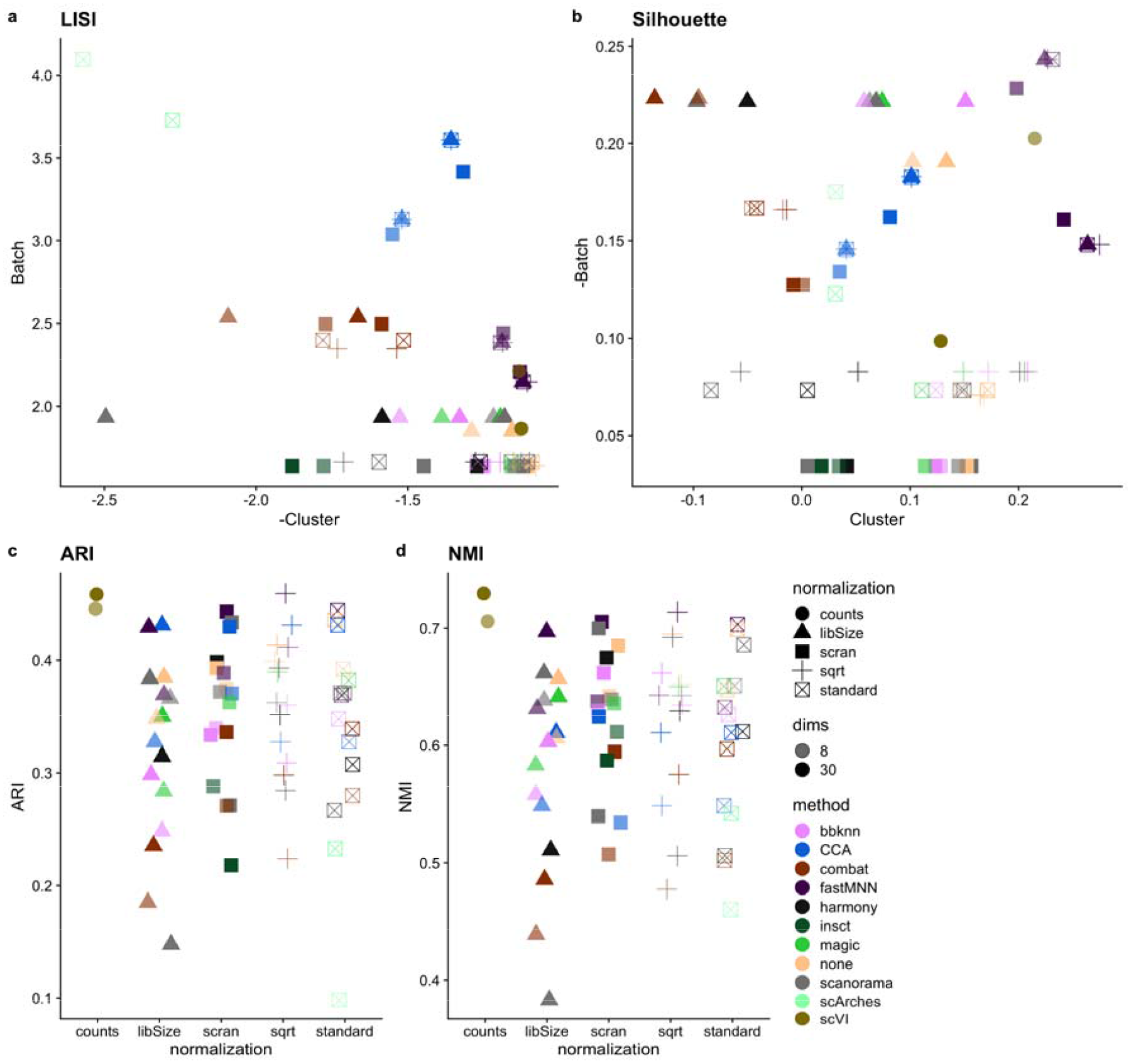
Performance of the various batch correction tools across various benchmarking metrics. For the LISI and Silhouette plots in A, B higher (y-axis) means better batch mixing and further to the right (x-axis) means better cluster purity. For the ARI and NMI metrics (which reflects how well cluster matches with cell type) in C, D, higher means a better score.

**Supplemental Figure 3:**
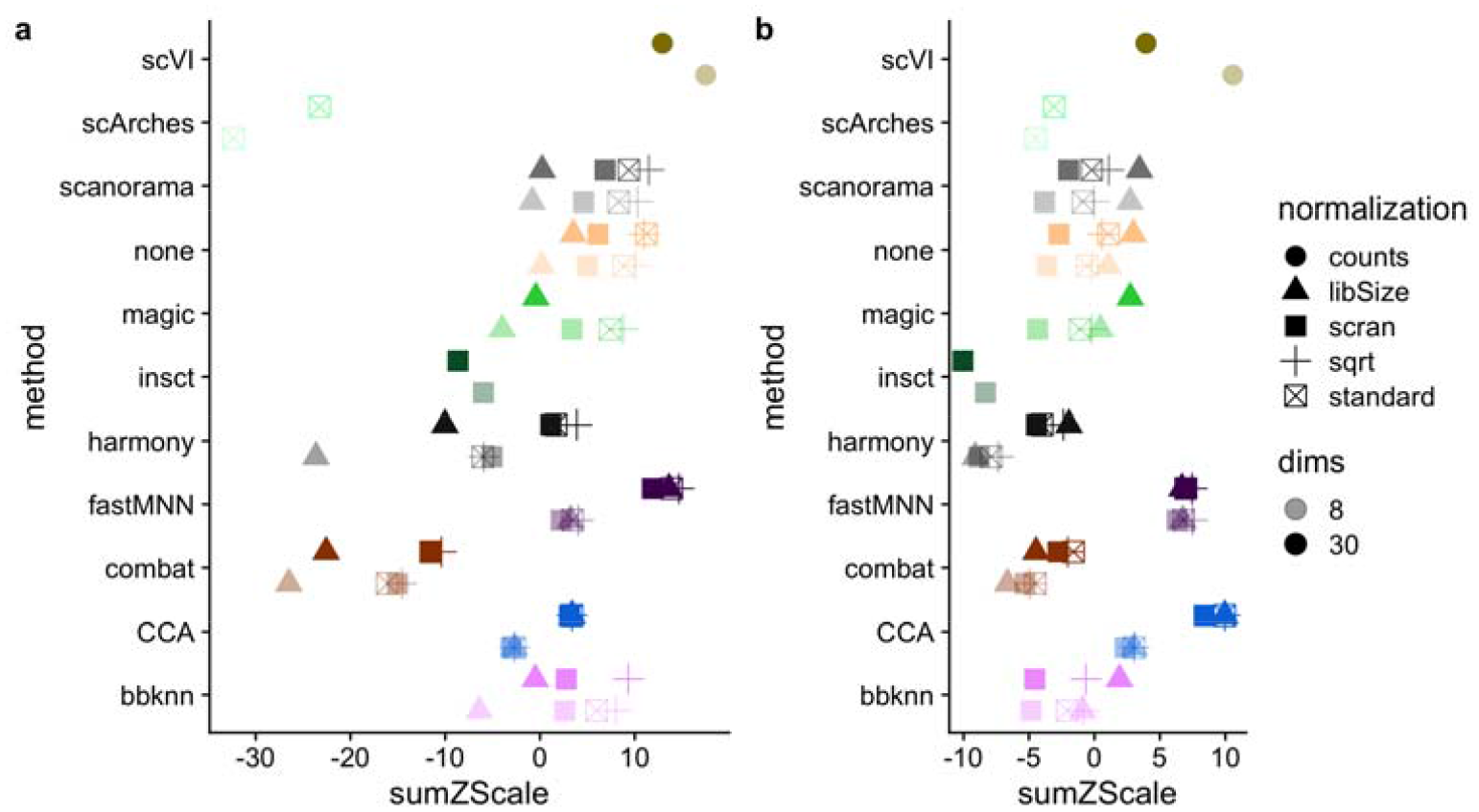
We take the sumZscoring from earlier, and apply two different weighting schemes to demonstrate how different priorities (much higher cluster purity or much more cluster mixing) can influence the scoring. In A we give a 3x multiplier to cell and cluster purity relative to batch mixing. In B we give a 3x multipler to batch mixing and we see that fastMNN with 8 dims has a higher score and the Seurat CCA metric ranks better. scVI performance with 8 latent dimension is consistently high across all weighting schemes.

**Supplemental Figure 4:**
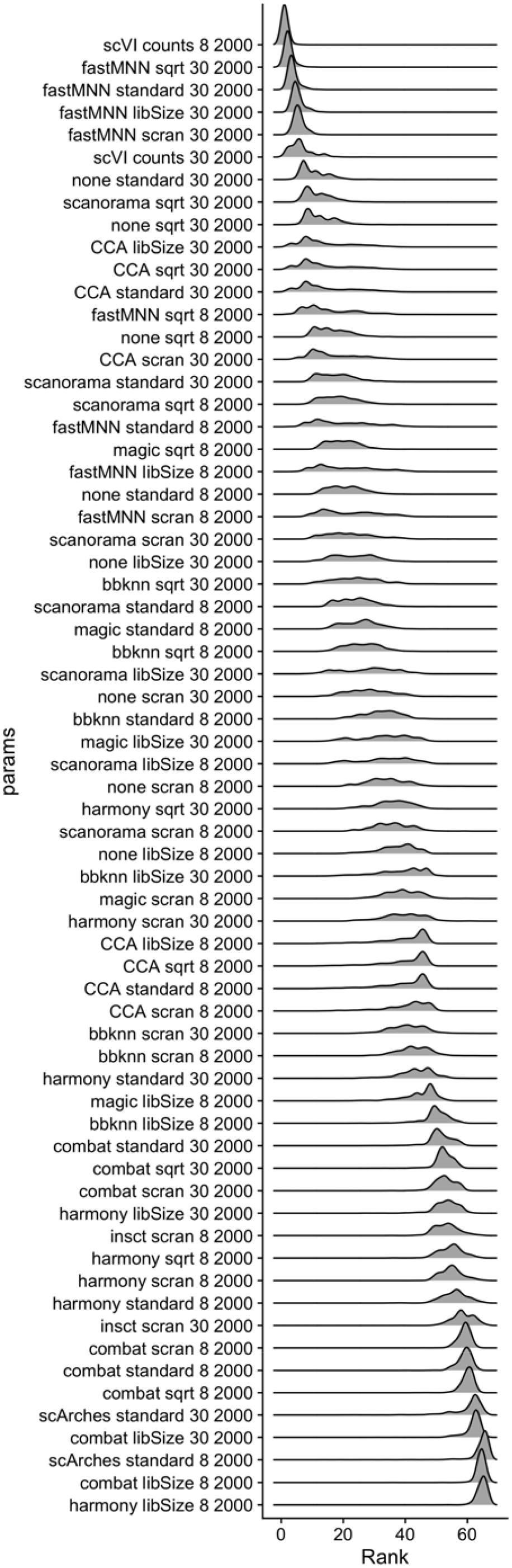
The sumZScale is composed of 10 metrics. We randomly weighed each zscaled metric by multiplying by a value randomly chosen between 0.1 and 10. This is bootstrapped 1000 times. The sumZScale is then computed and we extract the rank (by highest sumZScale) for 1000 bootstraps and plot the distribution as a density plot (lower rank is better integration performance). The y axis is ordered, top to bottom, by mean sumZScale Rank (higher on the y axis is better).

**Supplemental Figure 5:**
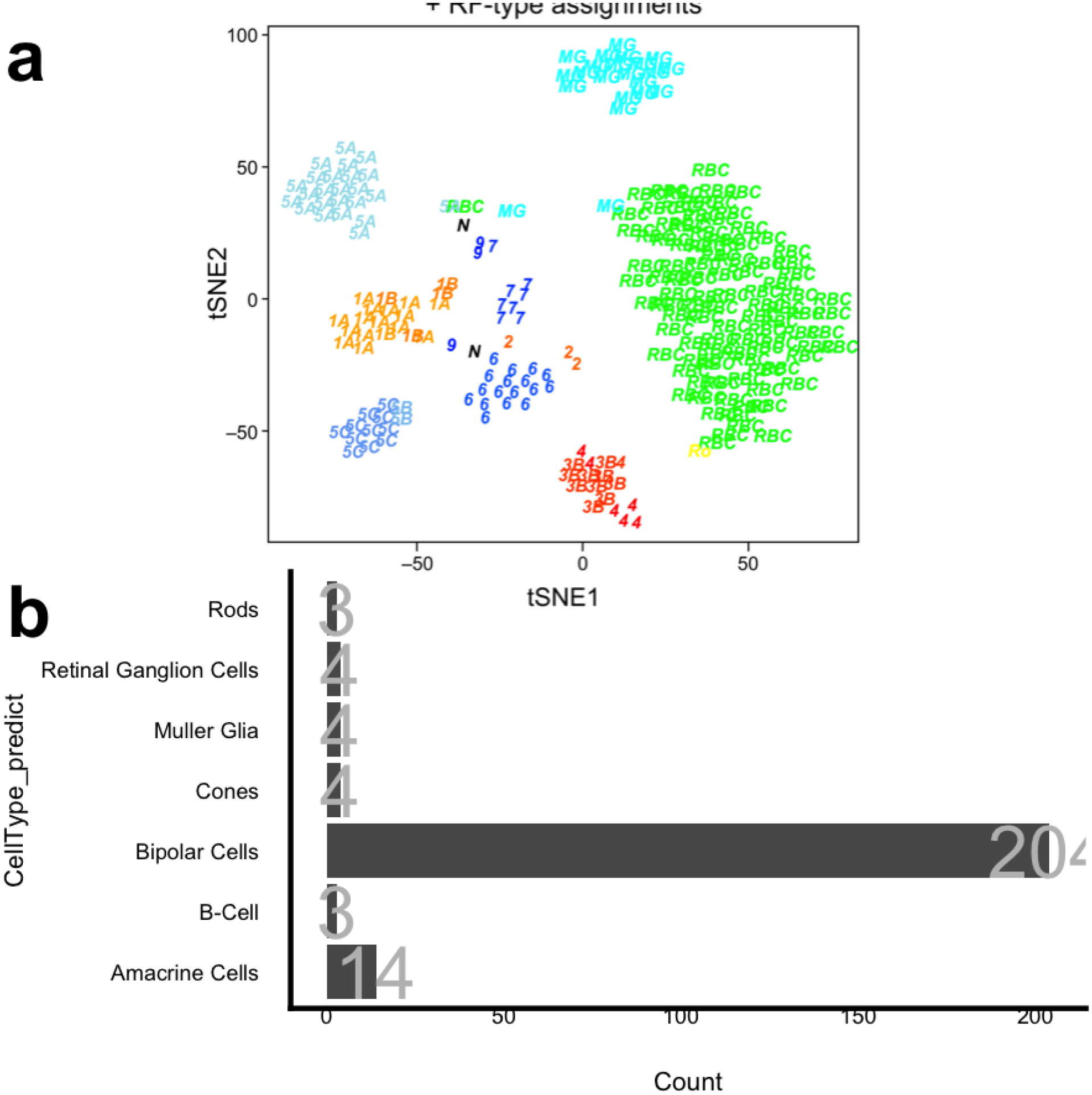
Our xgboost ML properly labels this Shekhar et al. RBC FAC sorted population as enriched in RBC

**Supplemental Figure 6:**
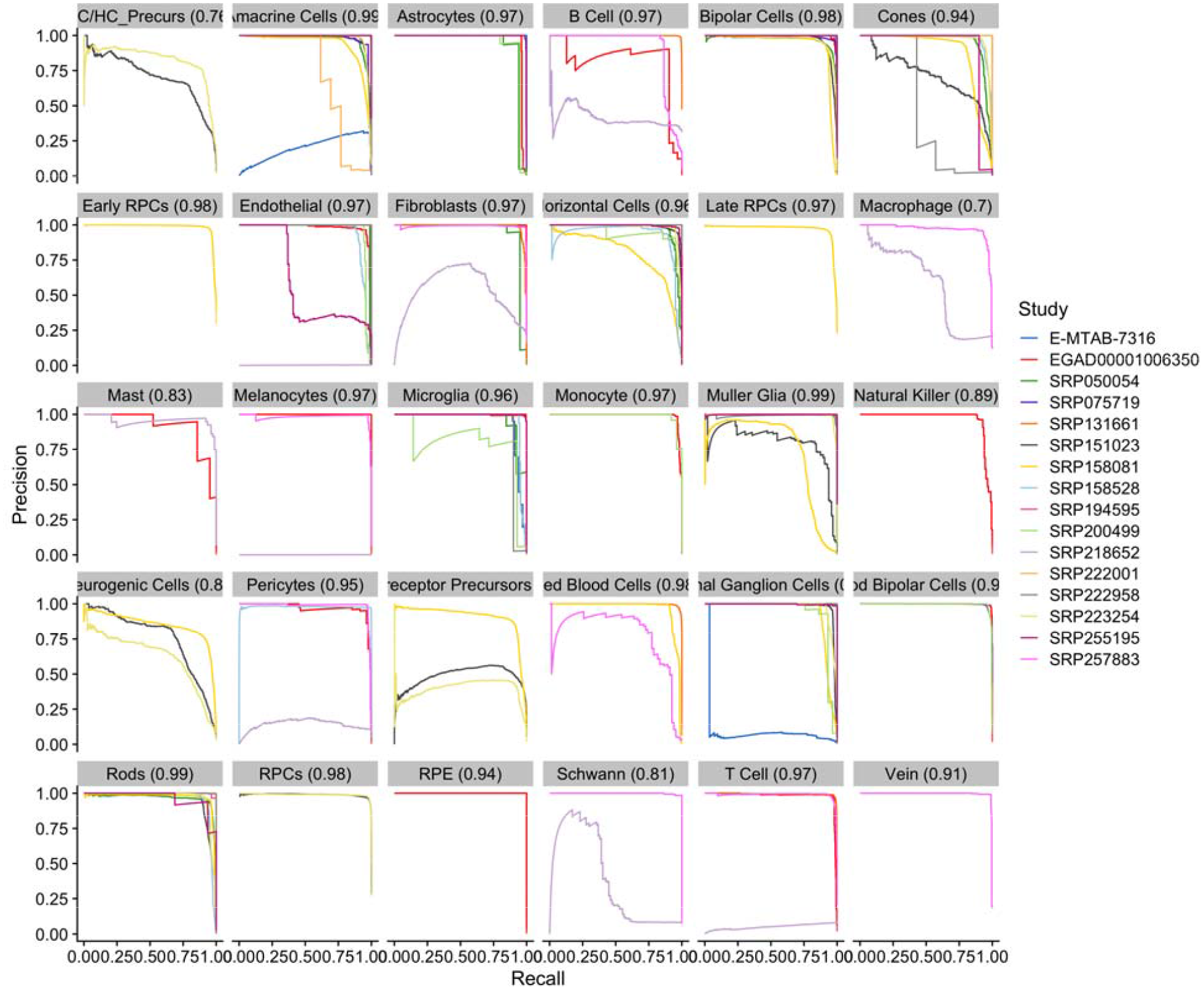
Precision recall curves for our xgboost cell type predictor model across each cell type predicted

**Supplemental Figure 7:**
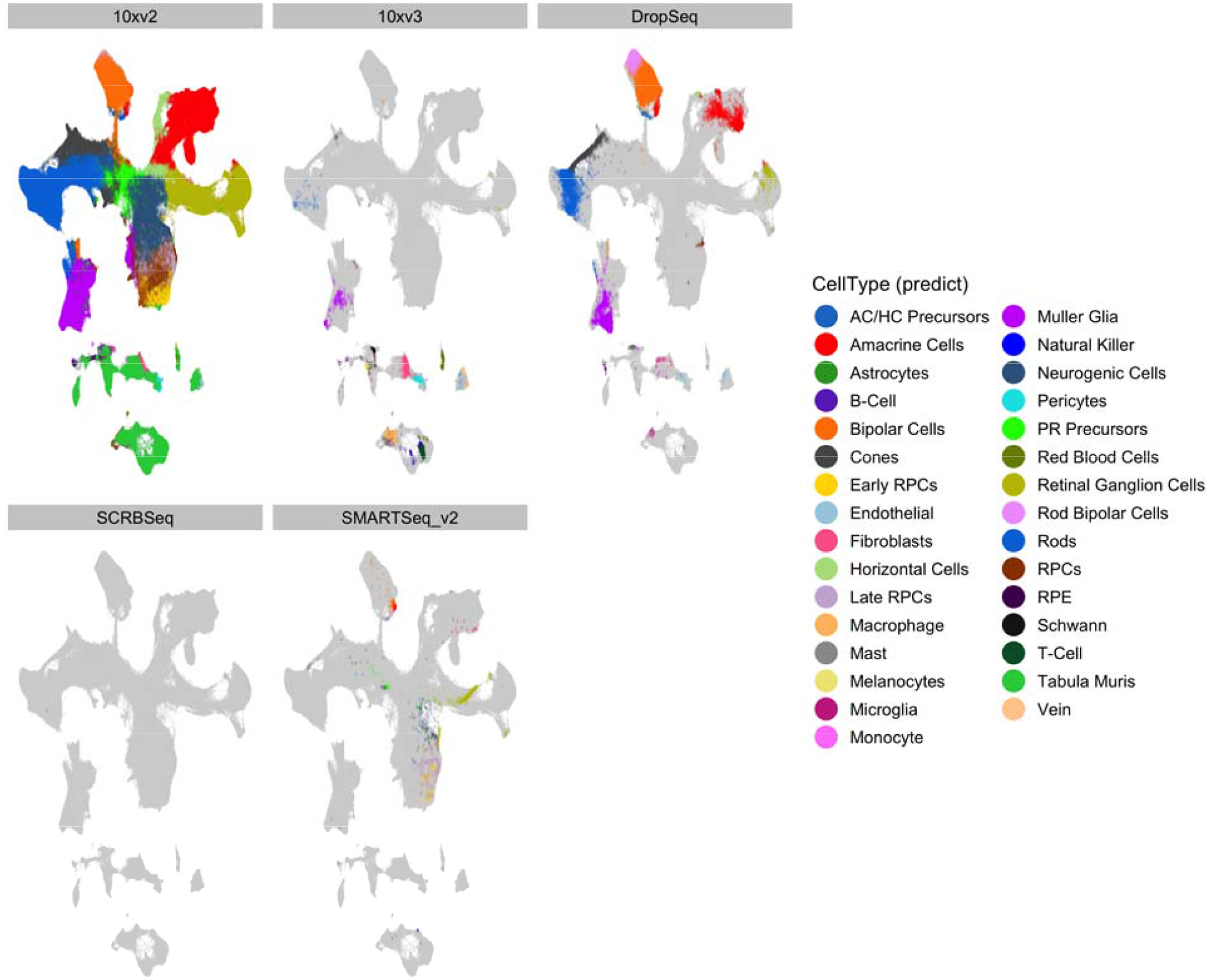
Distribution of cell types across the 2D UMAP confirms that the cell types are being properly placed despite the wide variety of single sequencing platforms present.

**Supplemental Figure 8:**
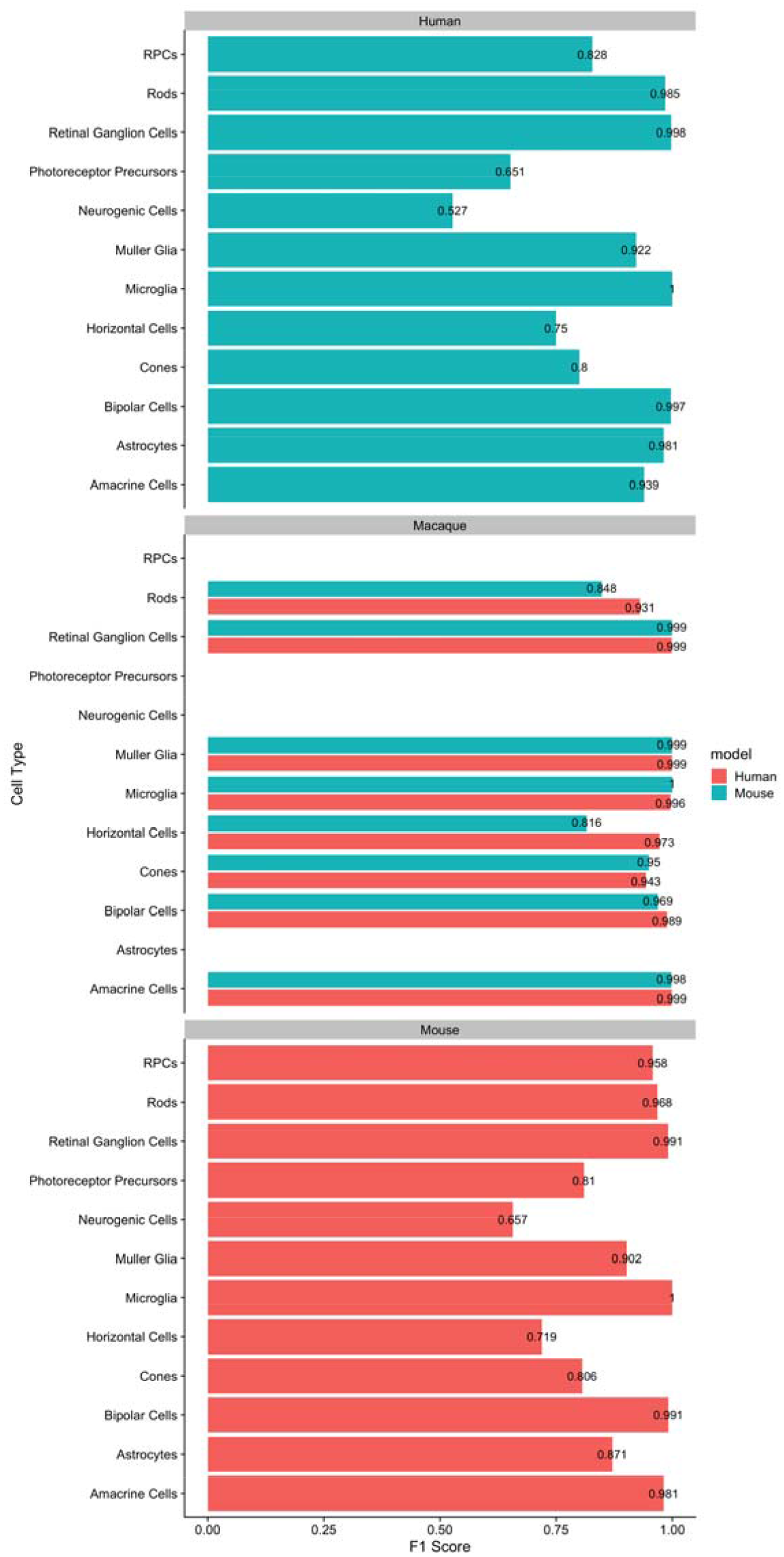
F1 scores (1 is perfect) of cell type prediction when using a human or mouse based xgboost cell type prediction model on other organisms (human, macaque, mouse).

**Supplemental Figure 9:**
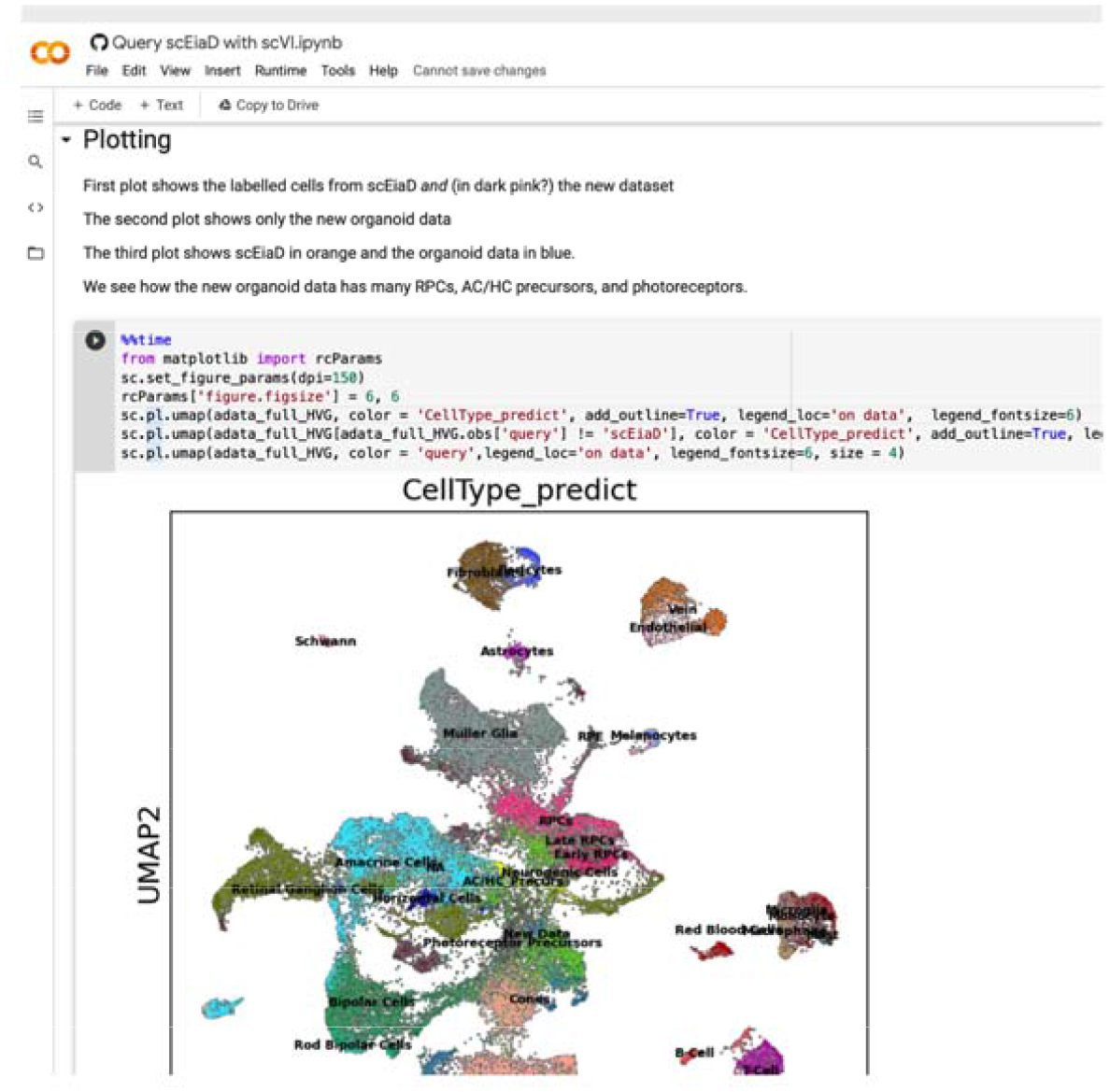
Screen shot of Google colab notebook that demonstrates how to integrate external data into the scEiaD resource. We see in the screenshot how the organoid dataset contains many of the retinal cell types.

**Supplemental Figure 10:**
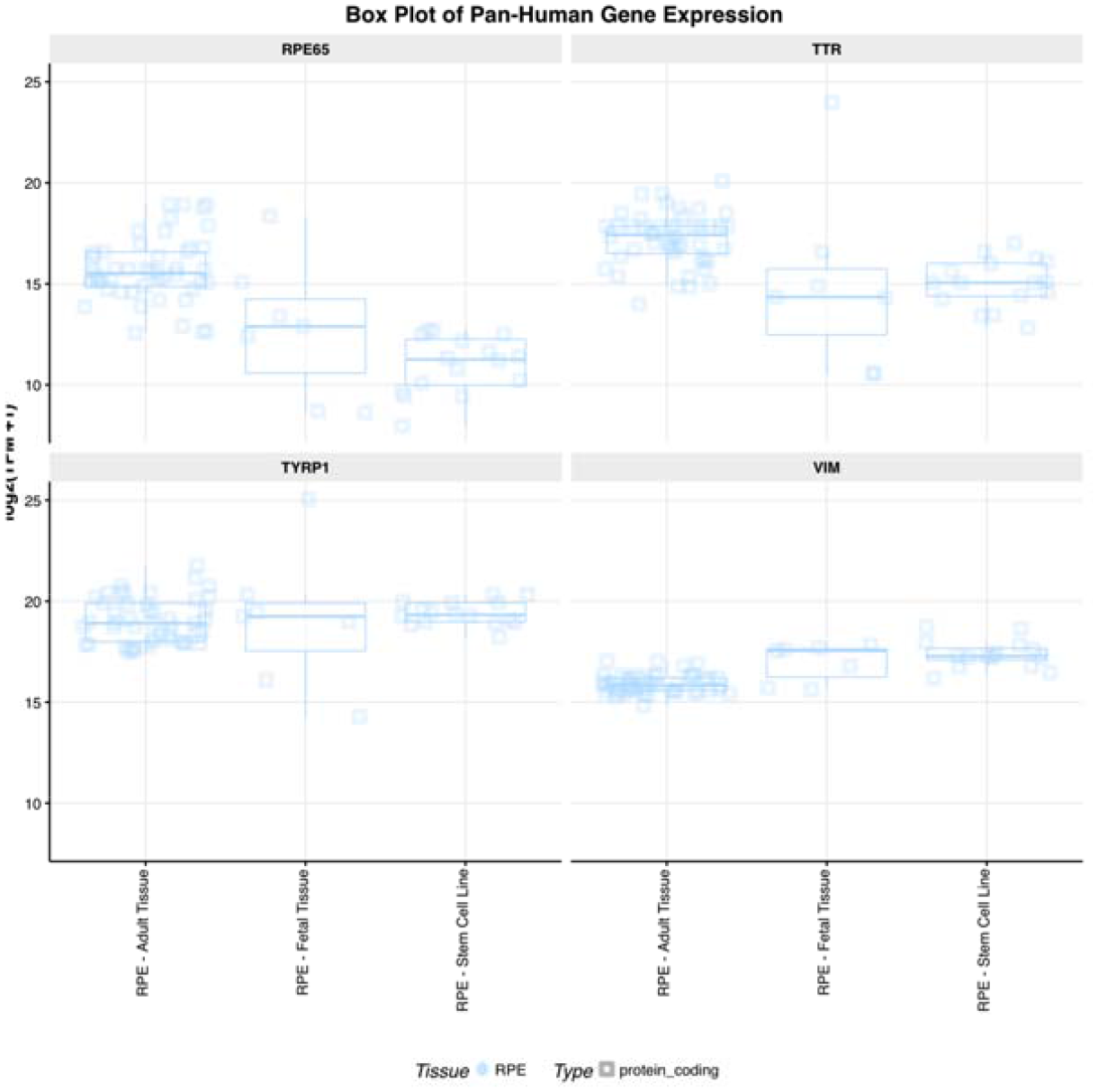
Screenshot of eyeIntegration bulk RNA-seq meta-analysis of vimentin expression in different RPE tissue sources

## Acknowledgments

We would like the thank the many groups who provided the raw data required to create this project. We keep a updated list of citations for the projects we pulled data from at https://plae.nei.nih.gov. We would also like to thank Adam Gayoso, who provided many useful comments on scVI parameter behavior and the Google colab implementation of scVI. Finally, this work utilized the computational resources of the NIH HPC Biowulf cluster (http://hpc.nih.gov).

## Works Cited

1. Masland RH. The Neuronal Organization of the Retina. Neuron. 2012;76(2):266–280. doi:10.1016/j.neuron.2012.10.002

2. Annies M, Kröger S. Isoform Pattern and AChR Aggregation Activity of Agrin Expressed by Embryonic Chick Retinal Ganglion Neurons. Molecular and Cellular Neuroscience. 2002;20(3):525–535. doi:10.1006/mcne.2002.1125

3. Hagstrom SA, Neitz M, Neitz J. Cone pigment gene expression in individual photoreceptors and the chromatic topography of the retina. J Opt Soc Am A Opt Image Sci Vis. 2000;17(3):527–537. doi:10.1364/josaa.17.000527

4. Trimarchi JM, Stadler MB, Roska B, et al. Molecular heterogeneity of developing retinal ganglion and amacrine cells revealed through single cell gene expression profiling. Journal of Comparative Neurology. 2007;502(6):1047–1065. doi:10.1002/cne.21368

5. Wahlin KJ, Lim L, Grice EA, Campochiaro PA, Zack DJ, Adler R. A method for analysis of gene expression in isolated mouse photoreceptor and Müller cells. Mol Vis. 2004;10:366–375.

6. Svensson V, da Veiga Beltrame E, Pachter L. A curated database reveals trends in single-cell transcriptomics. Database. 2020;2020. doi:10.1093/database/baaa073

7. Macosko EZ, Basu A, Satija R, et al. Highly Parallel Genome-wide Expression Profiling of Individual Cells Using Nanoliter Droplets. Cell. 2015;161(5):1202–1214. doi:10.1016/j.cell.2015.05.002

8. Buenaventura DF, Corseri A, Emerson MM. Identification of Genes With Enriched Expression in Early Developing Mouse Cone Photoreceptors. Invest Ophthalmol Vis Sci. 2019;60(8):2787–2799. doi:10.1167/iovs.19-26951

9. Clark BS, Stein-O’Brien GL, Shiau F, et al. Single-Cell RNA-Seq Analysis of Retinal Development Identifies NFI Factors as Regulating Mitotic Exit and Late-Born Cell Specification. Neuron. Published online May 22, 2019. doi:10.1016/j.neuron.2019.04.010

10. Cowan CS, Renner M, De Gennaro M, et al. Cell Types of the Human Retina and Its Organoids at Single-Cell Resolution. Cell. 2020;182(6):1623–1640.e34. doi:10.1016/j.cell.2020.08.013

11. Dharmat R, Kim S, Liu H, Fu S, Li Y, Chen R. Epigenetic adaptation prolongs photoreceptor survival during retinal degeneration. bioRxiv. Published online September 18, 2019:774950. doi:10.1101/774950

12. Fadl BR, Brodie SA, Malasky M, et al. An optimized protocol for retina single-cell RNA sequencing. Mol Vis. 2020;26:705–717. Accessed March 26, 2021. https://www.ncbi.nlm.nih.gov/pmc/articles/PMC7553720/

13. Hu Y, Wang X, Hu B, et al. Dissecting the transcriptome landscape of the human fetal neural retina and retinal pigment epithelium by single-cell RNA-seq analysis. PLoS Biol. 2019;17(7):e3000365. doi:10.1371/journal.pbio.3000365

14. Lehmann GL, Hanke-Gogokhia C, Hu Y, et al. Single-cell profiling reveals an endothelium-mediated immunomodulatory pathway in the eye choroid. Journal of Experimental Medicine. 2020;217(e20190730). doi:10.1084/jem.20190730

15. Lo Giudice Q, Leleu M, La Manno G, Fabre PJ. Single-cell transcriptional logic of cell-fate specification and axon guidance in early-born retinal neurons. Development. 2019;146(17). doi:10.1242/dev.178103

16. Lukowski SW, Lo CY, Sharov AA, et al. A single-cell transcriptome atlas of the adult human retina. EMBO J. 2019;38(18):e100811. doi:10.15252/embj.2018100811

17. Lu Y, Shiau F, Yi W, et al. Single-Cell Analysis of Human Retina Identifies Evolutionarily Conserved and Species-Specific Mechanisms Controlling Development. Dev Cell. 2020;53(4):473–491.e9. doi:10.1016/j.devcel.2020.04.009

18. Menon M, Mohammadi S, Davila-Velderrain J, et al. Single-cell transcriptomic atlas of the human retina identifies cell types associated with age-related macular degeneration. Nat Commun. 2019;10(1):4902. doi:10.1038/s41467-019-12780-8

19. O’Koren EG, Yu C, Klingeborn M, et al. Microglial Function Is Distinct in Different Anatomical Locations during Retinal Homeostasis and Degeneration. Immunity. 2019;50(3):723–737.e7. doi:10.1016/j.immuni.2019.02.007

20. Peng Y-R, Shekhar K, Yan W, et al. Molecular Classification and Comparative Taxonomics of Foveal and Peripheral Cells in Primate Retina. Cell. 2019;176(5):1222–1237.e22. doi:10.1016/j.cell.2019.01.004

21. Shekhar K, Lapan SW, Whitney IE, et al. Comprehensive Classification of Retinal Bipolar Neurons by Single-Cell Transcriptomics. Cell. 2016;166(5):1308–1323.e30. doi:10.1016/j.cell.2016.07.054

22. Sridhar A, Hoshino A, Finkbeiner CR, et al. Single-Cell Transcriptomic Comparison of Human Fetal Retina, hPSC-Derived Retinal Organoids, and Long-Term Retinal Cultures. Cell Rep. 2020;30(5):1644–1659.e4. doi:10.1016/j.celrep.2020.01.007

23. Tran NM, Shekhar K, Whitney IE, et al. Single-Cell Profiles of Retinal Ganglion Cells Differing in Resilience to Injury Reveal Neuroprotective Genes. Neuron. 2019;104(6):1039–1055.e12. doi:10.1016/j.neuron.2019.11.006

24. Voigt AP, Whitmore SS, Mulfaul K, et al. Bulk and single-cell gene expression analyses reveal aging human choriocapillaris has pro-inflammatory phenotype. Microvascular Research. 2020;131:104031. doi:10.1016/j.mvr.2020.104031

25. Voigt AP, Whitmore SS, Flamme-Wiese MJ, et al. Molecular characterization of foveal versus peripheral human retina by single-cell RNA sequencing. Experimental Eye Research. 2019;184:234–242. doi:10.1016/j.exer.2019.05.001

26. Voigt AP, Binkley E, Flamme-Wiese MJ, et al. Single-Cell RNA Sequencing in Human Retinal Degeneration Reveals Distinct Glial Cell Populations. Cells. 2020;9(2). doi:10.3390/cells9020438

27. Voigt AP, Mulfaul K, Mullin NK, et al. Single-cell transcriptomics of the human retinal pigment epithelium and choroid in health and macular degeneration. Proc Natl Acad Sci U S A. 2019;116(48):24100–24107. doi:10.1073/pnas.1914143116

28. Yan W, Peng Y-R, van Zyl T, et al. Cell Atlas of The Human Fovea and Peripheral Retina. Sci Rep. 2020;10(1):9802. doi:10.1038/s41598-020-66092-9

29. Yan W, Laboulaye MA, Tran NM, Whitney IE, Benhar I, Sanes JR. Mouse Retinal Cell Atlas: Molecular Identification of over Sixty Amacrine Cell Types. J Neurosci. 2020;40(27):5177–5195. doi:10.1523/JNEUROSCI.0471-20.2020

30. Melsted P, Booeshaghi AS, Gao F, et al. Modular and efficient pre-processing of single-cell RNA-seq. bioRxiv. Published online July 26, 2019:673285. doi:10.1101/673285

31. Srivastava A, Malik L, Smith T, Sudbery I, Patro R. Alevin efficiently estimates accurate gene abundances from dscRNA-seq data. Genome Biology. 2019;20(1):65. doi:10.1186/s13059-019-1670-y

32. Butler A, Hoffman P, Smibert P, Papalexi E, Satija R. Integrating single-cell transcriptomic data across different conditions, technologies, and species. Nature Biotechnology. 2018;36(5, 5):411–420. doi:10.1038/nbt.4096

33. Haghverdi L, Lun ATL, Morgan MD, Marioni JC. Batch effects in single-cell RNA-sequencing data are corrected by matching mutual nearest neighbors. Nature Biotechnology. 2018;36(5, 5):421–427. doi:10.1038/nbt.4091

34. Hie B, Bryson B, Berger B. Efficient integration of heterogeneous single-cell transcriptomes using Scanorama. Nature Biotechnology. 2019;37(6, 6):685–691. doi:10.1038/s41587-019-0113-3

35. Johnson WE, Li C, Rabinovic A. Adjusting batch effects in microarray expression data using empirical Bayes methods. Biostatistics. 2007;8(1):118–127. doi:10.1093/biostatistics/kxj037

36. Korsunsky I, Millard N, Fan J, et al. Fast, sensitive and accurate integration of single-cell data with Harmony. Nature Methods. 2019;16(12, 12):1289–1296. doi:10.1038/s41592-019-0619-0

37. Liu J, Gao C, Sodicoff J, Kozareva V, Macosko EZ, Welch JD. Jointly defining cell types from multiple single-cell datasets using LIGER. Nature Protocols. 2020;15(11, 11):3632–3662. doi:10.1038/s41596-020-0391-8

38. Lopez R, Regier J, Cole MB, Jordan MI, Yosef N. Deep generative modeling for single-cell transcriptomics. Nature Methods. 2018;15(12, 12):1053–1058. doi:10.1038/s41592-018-0229-2

39. Polański K, Young MD, Miao Z, Meyer KB, Teichmann SA, Park J-E. BBKNN: Fast batch alignment of single cell transcriptomes. Bioinformatics. 2020;36(3):964–965. doi:10.1093/bioinformatics/btz625

40. Query to reference single-cell integration with transfer learning | bioRxiv. Accessed December 20, 2020. https://www.biorxiv.org/content/10.1101/2020.07.16.205997v1

41. Simon LM, Wang Y-Y, Zhao Z. INSCT: Integrating millions of single cells using batch-aware triplet neural networks. bioRxiv. Published online May 17, 2020:2020.05.16.100024. doi:10.1101/2020.05.16.100024

42. Stuart T, Butler A, Hoffman P, et al. Comprehensive Integration of Single-Cell Data. Cell. 2019;177(7):1888–1902.e21. doi:10.1016/j.cell.2019.05.031

43. van Dijk D, Sharma R, Nainys J, et al. Recovering Gene Interactions from Single-Cell Data Using Data Diffusion. Cell. 2018;174(3):716–729.e27. doi:10.1016/j.cell.2018.05.061

44. Luecken MD, Büttner M, Chaichoompu K, et al. Benchmarking atlas-level data integration in single-cell genomics. bioRxiv. Published online May 27, 2020:2020.05.22.111161. doi:10.1101/2020.05.22.111161

45. Tran HTN, Ang KS, Chevrier M, et al. A benchmark of batch-effect correction methods for single-cell RNA sequencing data. Genome Biology. 2020;21(1):12. doi:10.1186/s13059-019-1850-9

46. Luecken MD, Theis FJ. Current best practices in single-cell RNA-seq analysis: A tutorial. Molecular Systems Biology. 2019;15(6):e8746. doi:10.15252/msb.20188746

47. Vandenbon A, Diez D. A clustering-independent method for finding differentially expressed genes in single-cell transcriptome data. Nature Communications. 2020;11(1, 1):4318. doi:10.1038/s41467-020-17900-3

48. Kallman A, Capowski EE, Wang J, et al. Investigating cone photoreceptor development using patient-derived NRL null retinal organoids. Commun Biol. 2020;3(1):82. doi:10.1038/s42003-020-0808-5

49. Hunt RC, Davis AA. Altered expression of keratin and vimentin in human retinal pigment epithelial cells in vivo and in vitro. J Cell Physiol. 1990;145(2):187–199. doi:10.1002/jcp.1041450202

50. Swamy V, McGaughey D. Eye in a Disk: eyeIntegration Human Pan-Eye and Body Transcriptome Database Version 1.0. Invest Ophthalmol Vis Sci. 2019;60(8):3236–3246. doi:10.1167/iovs.19-27106

51. Köster J, Rahmann S. Snakemake—a scalable bioinformatics workflow engine. Bioinformatics. 2012;28(19):2520–2522. doi:10.1093/bioinformatics/bts480

52. Harrow J, Frankish A, Gonzalez JM, et al. GENCODE: The reference human genome annotation for The ENCODE Project. Genome Res. 2012;22(9):1760–1774. doi:10.1101/gr.135350.111

53. J H, A F, Jm G, et al. GENCODE: The reference human genome annotation for The ENCODE Project. Genome Res. 2012;22(9):1760–1774. doi:10.1101/gr.135350.111

54. Yates AD, Achuthan P, Akanni W, et al. Ensembl 2020. Nucleic Acids Research. 2020;48(D1):D682–D688. doi:10.1093/nar/gkz966

55. Bray NL, Pimentel H, Melsted P, Pachter L. Near-optimal probabilistic RNA-seq quantification. Nat Biotechnol. 2016;34(5):525–527. doi:10.1038/nbt.3519

56. Melsted P, Ntranos V, Pachter L. The barcode, UMI, set format and BUStools. Bioinformatics. 2019;35(21):4472–4473. doi:10.1093/bioinformatics/btz279

57. Lun ATL, Riesenfeld S, Andrews T, et al. EmptyDrops: Distinguishing cells from empty droplets in droplet-based single-cell RNA sequencing data. Genome Biology. 2019;20(1):63. doi:10.1186/s13059-019-1662-y

58. Lun ATL, McCarthy DJ, Marioni JC. A step-by-step workflow for low-level analysis of single-cell RNA-seq data with Bioconductor. F1000Res. 2016;5:2122. doi:10.12688/f1000research.9501.2

59. Blondel VD, Guillaume J-L, Lambiotte R, Lefebvre E. Fast unfolding of communities in large networks. J Stat Mech. 2008;2008(10):P10008. doi:10.1088/1742-5468/2008/10/P10008

60. Traag V, Waltman L, van Eck NJ. From Louvain to Leiden: Guaranteeing well-connected communities. Sci Rep. 2019;9(1):5233. doi:10.1038/s41598-019-41695-z

61. Stassen SV, Siu DMD, Lee KCM, Ho JWK, So HKH, Tsia KK. PARC: Ultrafast and accurate clustering of phenotypic data of millions of single cells. Bioinformatics. 2020;36(9):2778–2786. doi:10.1093/bioinformatics/btaa042

62. McInnes L, Healy J, Melville J. UMAP: Uniform Manifold Approximation and Projection for Dimension Reduction. Published September 17, 2020. Accessed December 21, 2020. http://arxiv.org/abs/1802.03426

63. Büttner M, Miao Z, Wolf FA, Teichmann SA, Theis FJ. A test metric for assessing single-cell RNA-seq batch correction. Nature Methods. 2019;16(1, 1):43–49. doi:10.1038/s41592-018-0254-1

64. Gayoso A, Shor J. JonathanShor/DoubletDetection: Doubletdetection V3.0. Zenodo; 2020. doi:10.5281/zenodo.4359992

65. Wolock SL, Lopez R, Klein AM. Scrublet: Computational Identification of Cell Doublets in Single-Cell Transcriptomic Data. Cell Systems. 2019;8(4):281–291.e9. doi:10.1016/j.cels.2018.11.005

66. Bergen V, Lange M, Peidli S, Wolf FA, Theis FJ. Generalizing RNA velocity to transient cell states through dynamical modeling. Nature Biotechnology. 2020;38(12, 12):1408–1414. doi:10.1038/s41587-020-0591-3

67. Probabilistic harmonization and annotation of single-cell transcriptomics data with deep generative models. Molecular Systems Biology. 2021;17(1):e9620. doi:10.15252/msb.20209620

