## Supplementary figures and images for "Building the Mega Single Cell Transcriptome Ocular Meta-Atlas"

### UMAP all methods, colored by cell type

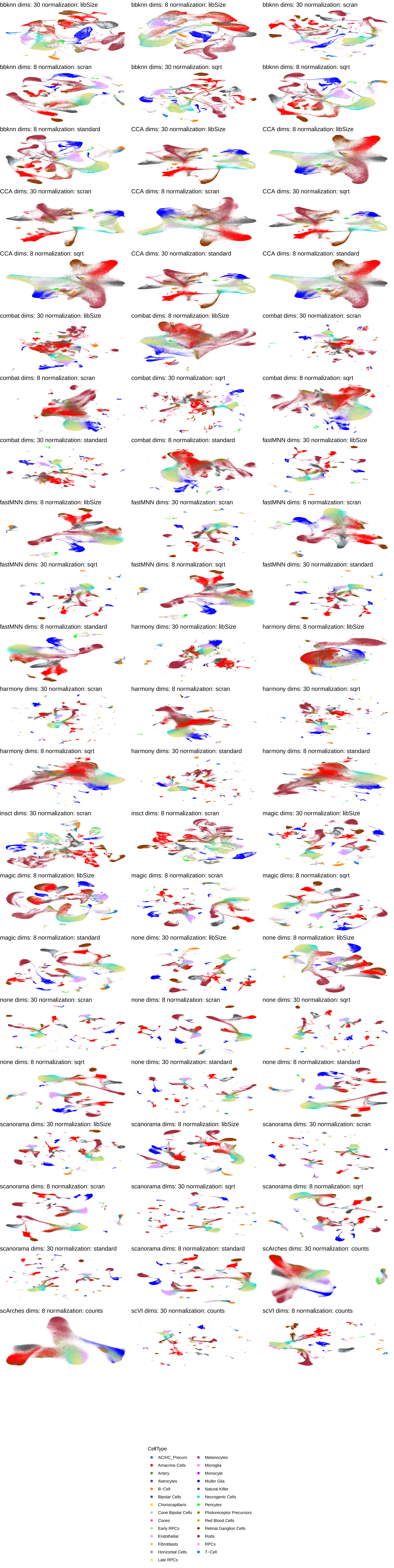

### UMAP all methods, colored by organism

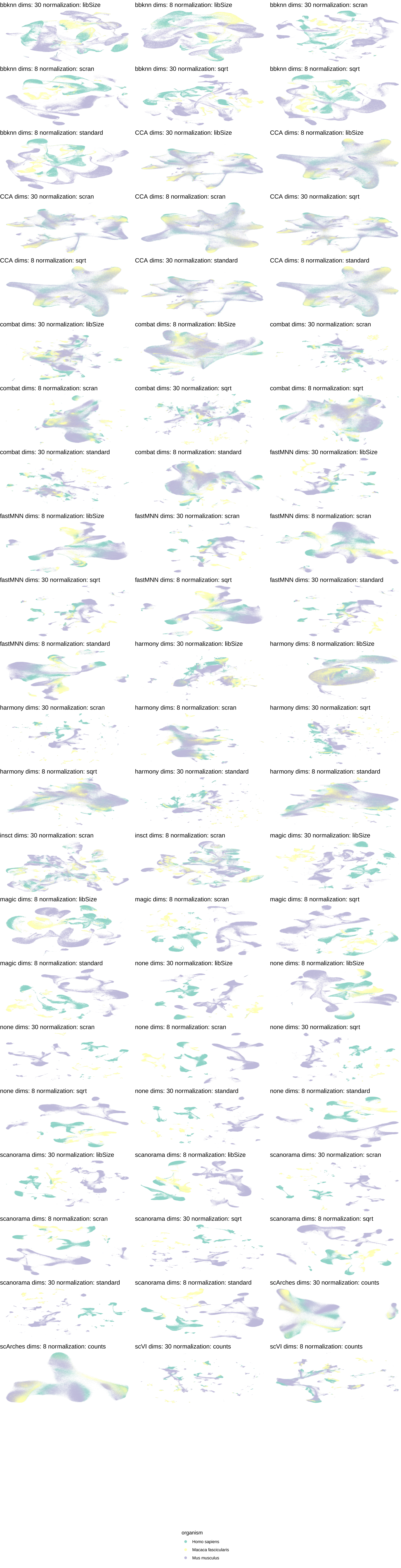

### UMAP all methods, colored by study accession

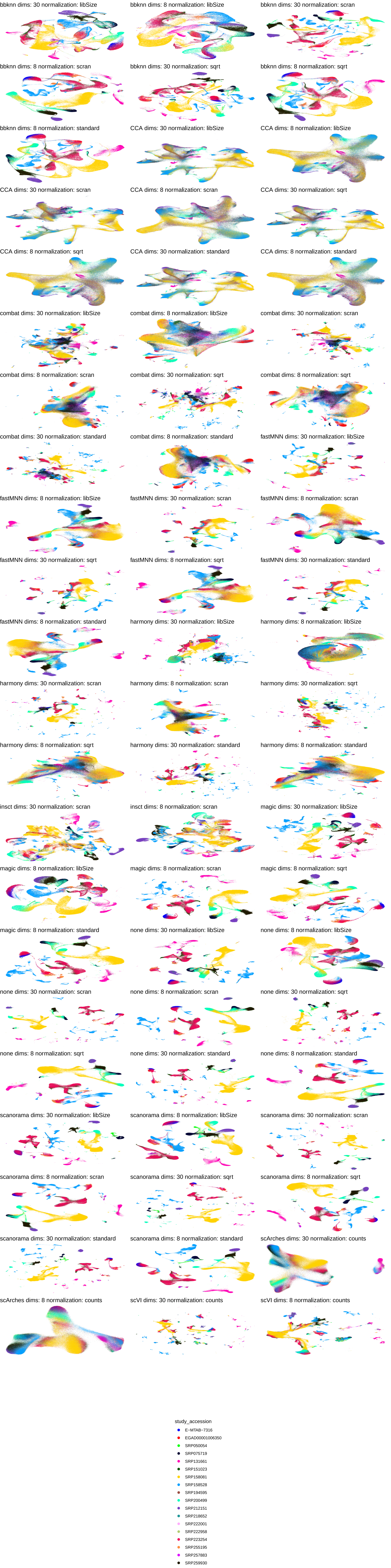
