## Supplementary material for "Building the Mega Single Cell Transcriptome Ocular Meta-Atlas": Seurat obj creation and integration implementation scripts: make_seurat_obj_functions.R

```
library(glue)
library(yaml)

make_seurat_obj <- function(m,
                            cell_info,
                            split.by = 'study_accession',
                            nfeatures = n_features,
                            keep_well = TRUE,
                            keep_droplet = TRUE,
                            mito_geneids,
							lengthCor = FALSE, 
							dont_use_well_for_FVF = FALSE,
							only_use_human_for_FVF = FALSE
                            ){
  well_m <- m[,cell_info %>% filter(value %in% colnames(m), !Platform %in% c('DropSeq', '10xv2', '10xv3')) %>% pull(value)]
  droplet_m <- m[,cell_info %>% filter(value %in% colnames(m), Platform %in% c('DropSeq', '10xv2', '10xv3')) %>% pull(value)]
  if (keep_well){
     seurat_well <- CreateSeuratObject(well_m)
  }
  if (keep_droplet){
     seurat_droplet <- CreateSeuratObject(droplet_m)
	 droplet_hs <- m[,cell_info %>% filter(value %in% colnames(m), organism %in% c('Homo sapiens')) %>% pull(value)]
	 hs_seurat_droplet <- CreateSeuratObject(droplet_hs)
  }

  # FILTER STEP!!!!
  # keep cells with < 10% mito genes, and more than 200 and less than 3000 detected genes for UMI
  # for well, drop the 3000 gene top end filter as there shouldn't be any droplets
  if (keep_well & !lengthCor){
    print('No Length Correction')
    seurat_well <- subset(seurat_well, subset = nFeature_RNA > 200)
  } else if (keep_well && lengthCor) {
      seurat_well <- subset(seurat_well, subset = nFeature_RNA > 200)
      #qumi_counts <- quminorm(seurat_well@assays$RNA@counts)
      source(glue('{git_dir}/src/extract_gene_length.R'))
    	geneL_mm <- gene_length( 'references/gtf/mm-mus_musculus_anno.gtf.gz')
    	geneL_hs <-  gene_length('references/gtf/hs-homo_sapiens_anno.gtf.gz')
    	well_hs <- cell_info %>% filter(Platform %in% c('C1', 'SCRBSeq', 'SMARTerSeq_v3', 'SMARTSeq_v2', 'SMARTSeq_v4'), organism == 'Homo sapiens') %>% pull(value)
      well_mm <- cell_info %>% filter(Platform %in% c('C1', 'SCRBSeq', 'SMARTerSeq_v3', 'SMARTSeq_v2', 'SMARTSeq_v4'), organism == 'Mus musculus') %>% pull(value)
    	mat <- seurat_well@assays$RNA@counts
    	hs_mat <- mat[, well_hs[well_hs %in% colnames(mat)]]
    	mm_mat <-  mat[, well_mm[well_mm %in% colnames(mat)]]
    	hs_mat_cor <- hs_mat / geneL_hs[row.names(hs_mat)] * median(geneL_hs)
    	mm_mat_cor <- mm_mat / geneL_mm[row.names(mm_mat)] * median(geneL_mm)
    	hs_mm <- cbind(hs_mat_cor, mm_mat_cor)
    	mat_cor <- hs_mm[, colnames(hs_mm) %in% colnames(mat)]

  	  seurat_well <- CreateSeuratObject(mat_cor)
      seurat_well <- subset(seurat_well, subset = nFeature_RNA > 200)
      print('Length Correction for Well DONE!')
  }
  # cells to keep
  if (!keep_well & keep_droplet){
    print('removing well')
    cells_to_keep <- row.names(seurat_droplet@meta.data)
  } else if (keep_well & !keep_droplet) {
    print('removing droplet')
    cells_to_keep <- row.names(seurat_well@meta.data)
  } else {
    print('keeping well and droplet')
    cells_to_keep <- c(row.names(seurat_droplet@meta.data), row.names(seurat_well@meta.data))
  }


  m_filter <- m[,cells_to_keep]


  seurat_m <- CreateSeuratObject(m_filter)
  mito_geneids_present <- mito_geneids[mito_geneids%in% rownames(seurat_m)]
  seurat_m[["percent.mt"]] <- PercentageFeatureSet(seurat_m, features = mito_geneids_present)
  seurat_m@meta.data$batch <- left_join(seurat_m@meta.data %>%
                                          row.names() %>% enframe(),
                                        cell_info, by = 'value') %>%
    pull(batch)
  seurat_m@meta.data$study_accession <- left_join(seurat_m@meta.data %>%
                                                    row.names() %>% enframe(),
                                                  cell_info, by = 'value') %>%
    pull(study_accession)
  seurat_m@meta.data$Age <- left_join(seurat_m@meta.data %>%
                                        row.names() %>% enframe(),
                                      cell_info, by = 'value') %>%
    pull(Age)
  seurat_m@meta.data$TechType <- left_join(seurat_m@meta.data %>%
                                        row.names() %>% enframe(),
                                      cell_info, by = 'value') %>%
    pull(Platform)

  ## account for any batches with less than 3 samples 
  batch_count <- table(seurat_m$batch) 
  bad_batches <- names(batch_count[batch_count<3])
  if(length(bad_batches) > 0){
    print('Removing the following batches which had ncells <3')
    print(bad_batches)
    good_cells <- filter(seurat_m@meta.data, !batch %in% bad_batches ) %>% rownames
    seurat_m <- seurat_m[,good_cells]
  }
  
  # scale data and regress
  seurat_m <- NormalizeData(seurat_m)
  # find var features
  seurat_m <- FindVariableFeatures(seurat_m, nfeatures = nfeatures, selection.method = 'vst')
  if (dont_use_well_for_FVF == TRUE) {
	seurat_mDrop <- FindVariableFeatures(seurat_droplet, nfeatures = nfeatures, selection.method = 'vst')
    VariableFeatures(seurat_m) <- VariableFeatures(seurat_mDrop)
  }

  if (only_use_human_for_FVF == TRUE) {
    hs_seurat_droplet  <- FindVariableFeatures(hs_seurat_droplet, nfeatures = nfeatures, selection.method = 'vst')
    VariableFeatures(seurat_m) <- VariableFeatures(hs_seurat_droplet)
  }  
  
  # don't use mito genes
  var_genes <- grep('^MT-', seurat_m@assays$RNA@var.features, value = TRUE, invert = TRUE)

  if (transform == 'standard'){
    print(paste0('Running lib.size and log correction, splitting by ', split.by))

    data <- seurat_m@assays$RNA@counts %>% t()
    data <- data[,var_genes]
    library_size <- Matrix::rowSums(data)
    median_transcript_count <- stats::median(library_size)
    data_norm <- median_transcript_count * data / library_size
    data_norm <- t(data_norm)
    data_norm <- log(data_norm + 1)
    seurat_m@assays$RNA@scale.data <- data_norm %>% as.matrix()
     #seurat_m <- ScaleData(seurat_m,
      #                    features = var_genes,
       #                   split.by = split.by,
        #                  do.center = TRUE,
         #                 do.scale = TRUE,
          #                verbose = TRUE,
           #               vars.to.regress = c("nCount_RNA", "nFeature_RNA", "percent.mt"))

    seurat_m <- RunPCA(seurat_m, npcs = 100)
  }
  seurat_m
}


# build SCT based seurat obj
seurat_sct <- function(seurat_list){
  ## tryCatch for SCT
  trySCTransform <- function(x){
    tryCatch(
      expr = {
        message("Successful SCTransform")
        return(SCTransform(x, vars.to.regress = c("nCount_RNA", "nFeature_RNA", "percent.mt")))
      },
      error = function(e){
        message("Failed SCTransform")
        print(e)
        return(NULL)
      }
    )
  }
  for (i in names(seurat_list)){
    DefaultAssay(seurat_list[[i]]) <- 'RNA'
    seurat_list[[i]] <- trySCTransform(seurat_list[[i]])
  }

  # remove sets with less than 500 cells, which will somehow(?) destory SCT - based integration performance
  low_n <- c()
  for (i in names(seurat_list)){
    if (ncol(seurat_list[[i]]) < 500){
      low_n <- c(low_n, i)
    }
  }
  if (length(low_n) > 0){
    seurat_list[low_n] <- NULL
  }

  study_data_features <- SelectIntegrationFeatures(object.list = seurat_list, nfeatures = n_features, verbose = FALSE)
  seurat_list <- PrepSCTIntegration(object.list = seurat_list, anchor.features = study_data_features, verbose = FALSE)

  # have to apply PCA after SelectIntegrationFeatures and PrepSCTIntegration
  seurat_list <- lapply(X = seurat_list, FUN = function(x) {
    x <- RunPCA(x, features = study_data_features, verbose = FALSE)
  })
  list(seurat_list = seurat_list, study_data_features = study_data_features)
}

#cran normalization
scran_norm <- function(seurat_obj = seurat__standard, split.by = 'batch'){
  var_features <- grep('^MT-', seurat_obj@assays$RNA@var.features, invert =TRUE, value = TRUE)
  seurat_list <- SplitObject(seurat_obj, split.by = covariate)
  print('Beginning scran norm')
  # list of seurat objects
  sce_list <- list()
  for (obj in names(seurat_list)){
    print(obj)
    print(seurat_list[[obj]] %>% dim())
    if (obj == 'SRP131661_10xv2_3-M-8'){
	  seurat_list[[obj]] <- NULL
	} else if (ncol(seurat_list[[obj]]) > 100){
      sce_list[[obj]] <- SingleCellExperiment(assays = list(counts = as.matrix(x = seurat_list[[obj]]$RNA@data)))
      clusters = quickCluster(sce_list[[obj]], min.size=100)
      sce_list[[obj]] = computeSumFactors(sce_list[[obj]], cluster=clusters)
      sce_list[[obj]] = scater::logNormCounts(sce_list[[obj]], log=TRUE)
      #sce_list[[obj]] = normalize(sce_list[[obj]], return_log = FALSE)
      print(summary(sizeFactors(sce_list[[obj]])))
      # seurat_list[[obj]]$RNA@data = as.sparse(log(x = assay(sce_list[[obj]], "normcounts") + 1))
      seurat_list[[obj]]$RNA@data = assay(sce_list[[obj]], "logcounts") %>% as.sparse()
    } else {seurat_list[[obj]] <- NULL} # remove obj with less than 50 cells
  }
  # merge back into one seurat obj
  merged <- merge(x = seurat_list[[1]], y = seurat_list[2:length(x = seurat_list)])
  merged$RNA@scale.data = merged$RNA@data[var_features,] %>% as.matrix()
  merged@assays$RNA@var.features <- var_features
  # re do PCA
  merged <- RunPCA(merged, npcs = 100, features = var_features)
  merged
}


#' Performs L1 normalization on input data such that the sum of expression
#' values for each cell sums to 1, then returns normalized matrix to the metric
#' space using median UMI count per cell effectively scaling all cells as if
#' they were sampled evenly.
#' @param seurat_object
#' @return seurat_obj
#' 2 dimensional array with normalized gene expression values
#' @import Matrix
#' @import dplyr
#'
#' @export
library.size.normalize <- function(seurat_obj, sqrt = FALSE, verbose=FALSE) {
  if (verbose) {
    message(paste0(
      "Normalizing library sizes for ",
      nrow(data), " cells"
    ))
  }
  vfeatures <- grep('^MT-', seurat_obj@assays$RNA@var.features, invert =TRUE, value = TRUE)
  data <- seurat_obj@assays$RNA@counts %>% t()
  data <- data[,vfeatures]
  library_size <- Matrix::rowSums(data)
  median_transcript_count <- stats::median(library_size)
  data_norm <- median_transcript_count * data / library_size
  data_norm <- t(data_norm)
  if (sqrt) {data_norm <- sqrt(data_norm)}
  seurat_obj@assays$RNA@scale.data <- data_norm %>% as.matrix()
  seurat_obj <- RunPCA(seurat_obj, features = vfeatures)
  seurat_obj
}
```
