## Supplementary material for "Building the Mega Single Cell Transcriptome Ocular Meta-Atlas": Seurat obj creation and integration implementation scripts: merge_methods.R

```
# run integration methods that support Seurat objects directly
args <- commandArgs(trailingOnly = TRUE)
system('mkdir -p loom')
method = args[1]

conda_dir = Sys.getenv('SCIAD_CONDA_DIR')
git_dir = Sys.getenv('SCIAD_GIT_DIR')
cat('\n\n')
cat(conda_dir)
cat('\n')
cat(git_dir)
cat('\n\n')
# crazy section to deal with that fact I have scanorama in a conda environment,
# but many of the seurat wrapped integration tools can't be installed in conda
# without crazy effort (e.g liger...as it needs to be compiled in C)
library(glue)
if (method == 'scanorama'){
  Sys.setenv(RETICULATE_PYTHON = glue('{conda_dir}/envs/scanorama/bin/python') )
  library(reticulate)
  use_condaenv("scanorama")
  scanorama <- import('scanorama')
  print('Scanorama imported')
} else if (method == 'magic') {
  Sys.setenv(RETICULATE_PYTHON = glue('{conda_dir}/envs/magic/bin/python') )
  library(reticulate)
  library(Rmagic)
} else {
  library(loomR)
  library(SeuratWrappers)
  library(harmony)
  library(batchelor)
  library(sva)
  library(Matrix)
}
library(data.table)
library(tidyverse)
library(Seurat)


transform = args[2]
covariate = args[3]
latent = args[4] %>% as.numeric()
#args[5] <- gsub('counts', 'standard', args[5])
load(args[5])


run_integration <- function(seurat_obj, method, covariate = 'study_accession', transform = 'standard', latent = 50, file = args[5], n_epochs = 5){
  # covariate MUST MATCH what was used in build_seurat_obj.R
  # otherwise weird-ness may happen
  # the scaling happens at this level
  # e.g. DO NOT use 'batch' in build_seurat_obj.R then 'study_accession' here
 if (method == 'bbknn'){
    if (transform == 'SCT'){
	  assay <- 'SCT'
      vfeatures <- grep('^MT-', seurat_obj@assays$SCT@var.features, invert =TRUE, value = TRUE)
      matrix = seurat_obj@assays$SCT@scale.data[vfeatures, ]     
	} else {
    assay <- 'RNA'
    vfeatures <- grep('^MT-', seurat_obj@assays$RNA@var.features, invert =TRUE, value = TRUE)
    matrix = seurat_obj@assays$RNA@scale.data[vfeatures, ]
	}
	rand <- sample(1e7:9e7,1)
    out <- paste0('loom/', method, '_', covariate, '_', transform, '_', length(vfeatures), '_', rand, '_',  latent, '.loom')
	create(filename= out, 
           overwrite = TRUE,
           data = matrix, 
           cell.attrs = list(batch = seurat_obj@meta.data[,covariate],
                             batch_indices = seurat_obj@meta.data[,covariate] %>% 
                               as.factor() %>% 
                               as.numeric()))
    # connect to new loom file, then disconnect...otherwise python call gets borked for 
    # as we are connected into the file on create
    loom <- connect(out, mode = 'r')
    loom$close_all() 
	

	bbknn_command = paste(glue('{conda_dir}/envs/bbknn/bin/./python {git_dir}/src/run_bbknn.py'), out, latent)
    # run bbknn 
    print(bbknn_command)
    system(bbknn_command)
	# import batch corrected dims	
    latent_dims <- read.csv(paste0(out, '.csv'), header = FALSE)
	latent_dims <- latent_dims[2:nrow(latent_dims),2:ncol(latent_dims)]
    row.names(latent_dims) <- colnames(seurat_obj)
    colnames(latent_dims) <- paste0("bbknn_", 1:ncol(latent_dims))
    
    seurat_obj[["bbknn"]] <- CreateDimReducObject(embeddings = latent_dims %>% as.matrix(), key = "bbknn_", assay = DefaultAssay(seurat_obj))
	obj <- seurat_obj 
 } 
 else if (method == 'CCA'){
    obj <- seurat_obj
    if (transform == 'SCT'){
      # remove sets with fewre than 1000 cells
      seurat_obj$seurat_list[seurat_obj$seurat_list %>% 
                               map(ncol) %>% 
                               map(enframe) %>%
                               bind_rows(.id = 'ID') %>% 
                               filter(value < 100) %>% 
                               pull(ID)] <- NULL
	  anchors <- FindIntegrationAnchors(object.list = seurat_obj$seurat_list, dims = 1:latent, normalization.method = 'SCT',
                                        anchor.features = seurat_obj$study_data_features) 
      obj <- IntegrateData(anchorset = anchors, verbose = TRUE, normalization.method = 'SCT')
    } else {
	  # downsample to 250k to make runtime not insane for gigascience resubmission
	  # at 4+ days right now I can't handle this shit anymore
	  
	  #obj <- seurat_obj[,sample(1:ncol(seurat_obj), size = 250000, replace = FALSE)]
      seurat_list <- SplitObject(obj, split.by = covariate)
      seurat_list[seurat_list %>% 
                    map(ncol) %>% 
                    map(enframe) %>%
                    bind_rows(.id = 'ID') %>% 
                    filter(value < 1000) %>% 
                    pull(ID)] <- NULL
      

      # RPCA
	  seurat_list <- lapply(X = seurat_list, FUN = function(x) {
         x <- ScaleData(x, features = obj[[obj@active.assay]]@var.features, verbose = FALSE)
         x <- RunPCA(x, features = obj[[obj@active.assay]]@var.features, verbose = FALSE)
      })
	  # integrate against the human data as it keeps exploding otherwise
	  # nightmare algorithm
      load('pipeline_data/cell_info/cell_info_labelled.Rdata')
	  refs <- cell_info_labels %>% select(organism, study_accession) %>% unique() %>% filter(organism == 'Homo sapiens', !grepl('OGV', study_accession)) %>% pull(study_accession)
      sref <- grep( paste(refs, collapse = "|"), names(seurat_list))
	  anchors <- FindIntegrationAnchors(object.list = seurat_list, dims = 1:latent, reduction = 'rpca', k.filter = NA, reference = sref,
                                        anchor.features = obj[[obj@active.assay]]@var.features)
						                #reduction = 'rpca')
      obj <- IntegrateData(anchorset = anchors, verbose = TRUE, dims = 1:latent)
    }
    DefaultAssay(obj) <- "integrated"
    obj <- ScaleData(obj)
    obj <- RunPCA(obj, npcs = 100)
  } else if (method == 'fastMNN'){
    ## uses list of seurat objects (each obj your "covariate")
    seurat_list <- SplitObject(seurat_obj, split.by = covariate)
    obj <- RunFastMNN(object.list = seurat_list, d = latent)
    # put back scaledata as it gets wiped
    if (transform != 'SCT'){
      var_genes <- grep('^MT-', seurat_obj@assays$RNA@var.features, value = TRUE, invert = TRUE)
      obj@assays$RNA@scale.data <- seurat_obj@assays$RNA@scale.data
    }
  } else if (method == 'magic') {
	print(transform)
	assay <- 'RNA'
    vfeatures <- grep('^MT-', seurat_obj@assays$RNA@var.features, invert =TRUE, value = TRUE)
    if (transform == 'sqrt'){
	  matrix = seurat_obj@assays$RNA@counts[vfeatures, ] %>% t()
      matrix = library.size.normalize(matrix)
      matrix = sqrt(matrix)
    } else if (transform == 'SCT'){
      assay <- 'SCT'
      vfeatures <- grep('^MT-', seurat_obj@assays$SCT@var.features, invert =TRUE, value = TRUE)
      matrix = seurat_obj@assays$SCT@scale.data[vfeatures, ] %>% Matrix(., sparse = TRUE) %>% t()
    } else {
      matrix = seurat_obj@assays$RNA@scale.data[vfeatures, ] %>% t()
    } 
	magic <- magic(matrix, genes = "pca_only", n.jobs=10, npca = latent, verbose = TRUE)
	seurat_obj[["magic"]] <- CreateDimReducObject(embeddings = 
								magic$result %>% as.matrix(), 
								key = "magic_", 
								assay = DefaultAssay(seurat_obj))
	obj <- seurat_obj
  } else if (method == 'insct') {
    assay <- 'RNA'
    vfeatures <- grep('^MT-', seurat_obj@assays$RNA@var.features, invert =TRUE, value = TRUE)
    if (transform == 'counts'){
      matrix = seurat_obj@assays$RNA@counts[vfeatures, ]
    } else if (transform == 'SCT'){
      assay <- 'SCT'
      vfeatures <- grep('^MT-', seurat_obj@assays$SCT@var.features, invert =TRUE, value = TRUE)
      matrix = seurat_obj@assays$SCT@scale.data[vfeatures, ] %>% Matrix(., sparse = TRUE)
    } else {
      matrix = seurat_obj@assays$RNA@scale.data[vfeatures, ]
    }
	if (ncol(matrix) < 100000){type = 'onlyWELL'} else {type = 'onlyDROPLET'}
    out <- paste0('loom/', method, '_', covariate, '_', transform, '_', length(vfeatures), '_', latent, '_', type, '.loom')
	
	# add count to one cell if all are zero
	vfeature_num <- length(vfeatures)
	one0 <- vector(mode = 'numeric', length = vfeature_num)
	one0[2] <- 1
	if (sum(colSums(matrix)==0) > 0){
		matrix[,colSums(matrix) == 0] <- one0
	}

    load('pipeline_data/cell_info/cell_info_labelled.Rdata')
    ct <- seurat_obj@meta.data %>% as_tibble(rownames = 'value') %>% left_join(cell_info_labels, by = 'value') %>% pull(CellType)
	ct[ct == 'Doublet'] <- 'Missing'
	ct[ct == 'Doublets'] <- 'Missing'
	ct[is.na(ct)] <- 'Missing'
	create(filename= out, 
           overwrite = TRUE,
           data = matrix, 
           cell.attrs = list(batch = seurat_obj@meta.data[,covariate],
							 masking_batch = cbind(ct, batch = seurat_obj@meta.data[,covariate]) %>% 
										as_tibble() %>% 
										mutate(mask_batch = case_when(ct == 'Missing' ~ 'missing', 
										 							  batch == 'SRP050054_DropSeq_retina5' ~ 'missing',
																	  batch == 'SRP050054_DropSeq_retina6' ~ 'missing',
																	  TRUE ~ batch)) %>% 
										pull(mask_batch),
							 celltype = ct,
                             batch_indices = seurat_obj@meta.data[,covariate] %>% 
                               as.factor() %>% 
                               as.numeric()))
    # connect to new loom file, then disconnect...otherwise python call gets borked for 
    # as we are connected into the file on create
    loom <- connect(out, mode = 'r')
    loom$close_all() 

    insct_command = paste(glue('{conda_dir}/envs/INSCT/bin/./python3.7 {git_dir}/src/run_INSCT.py'),
                         out, latent)
    # run insct     
	print(insct_command) 
    system(insct_command)
    # import reduced dim (latent)
    latent_dims <- read.csv(paste0(out, '.csv'), header = FALSE)
	latent_dims <- latent_dims[2:nrow(latent_dims),2:ncol(latent_dims)]
    row.names(latent_dims) <- colnames(seurat_obj)
    colnames(latent_dims) <- paste0("insct_", 1:ncol(latent_dims))
    
    seurat_obj[["insct"]] <- CreateDimReducObject(embeddings = latent_dims %>% as.matrix(), key = "insct_", assay = DefaultAssay(seurat_obj))
	obj <- seurat_obj 

  } else if (method == 'desc') {
    assay <- 'RNA'
    vfeatures <- grep('^MT-', seurat_obj@assays$RNA@var.features, invert =TRUE, value = TRUE)
    if (transform == 'counts'){
      matrix = seurat_obj@assays$RNA@counts[vfeatures, ]
    } else if (transform == 'SCT'){
      assay <- 'SCT'
      vfeatures <- grep('^MT-', seurat_obj@assays$SCT@var.features, invert =TRUE, value = TRUE)
      matrix = seurat_obj@assays$SCT@scale.data[vfeatures, ] %>% Matrix(., sparse = TRUE)
    } else {
      matrix = seurat_obj@assays$RNA@scale.data[vfeatures, ]
    }
	out <- paste0('loom/', method, '_', covariate, '_', transform, '_', length(vfeatures), '_', latent, '_', sample(1e5:5e5, 1), '.loom')
		
	# add count to one cell if all are zero
	vfeature_num <- length(vfeatures)
	one0 <- vector(mode = 'numeric', length = vfeature_num)
	one0[2] <- 1
	if (sum(colSums(matrix)==0) > 0){
		matrix[,colSums(matrix) == 0] <- one0
	}
	# desc is crapping out and dropping a features
	# remove the lowest 4 expression feature (genes)
	min_val = min_val = rowSums(matrix) %>% sort() %>% head(5) %>% tail(1)
	removal = (rowSums(matrix) < min_val)
	matrix <- matrix[!removal,]
	create(filename= out, 
           overwrite = TRUE,
           data = matrix, 
           cell.attrs = list(batch = seurat_obj@meta.data[,covariate],
                             batch_indices = seurat_obj@meta.data[,covariate] %>% 
                               as.factor() %>% 
                               as.numeric()))
    # connect to new loom file, then disconnect...otherwise python call gets borked for 
    # as we are connected into the file on create
    loom <- connect(out, mode = 'r')
    loom$close_all() 

    desc_command = paste(glue('{conda_dir}/envs/DESC/bin/./python3.7 {git_dir}/src/run_desc.py'),
                         out, transform, latent)
    # run desc     
	print(desc_command) 
    system(desc_command)
    # import reduced dim (latent)
    latent_dims <- read.csv(paste0(out, '.csv'), header = FALSE)
	latent_dims <- latent_dims[2:nrow(latent_dims),2:ncol(latent_dims)]
    row.names(latent_dims) <- colnames(seurat_obj)
    colnames(latent_dims) <- paste0("desc_", 1:ncol(latent_dims))
    
    seurat_obj[["desc"]] <- CreateDimReducObject(embeddings = latent_dims %>% as.matrix(), key = "desc_", assay = DefaultAssay(seurat_obj))
	obj <- seurat_obj 

  } else if (method == 'scArches') {
    # scArches ----
    assay <- 'RNA'
    vfeatures <- grep('^MT-', seurat_obj@assays$RNA@var.features, invert =TRUE, value = TRUE)
    if (transform == 'counts'){
      matrix = seurat_obj@assays$RNA@counts #[vfeatures, ]
    } else if (transform == 'SCT'){
      assay <- 'SCT'
      vfeatures <- grep('^MT-', seurat_obj@assays$SCT@var.features, invert =TRUE, value = TRUE)
      matrix = seurat_obj@assays$SCT@scale.data[vfeatures, ] %>% Matrix(., sparse = TRUE)
    } else {
      matrix = seurat_obj@assays$RNA@scale.data[vfeatures, ]
    }
	rand <- sample(1e7:9e7,1)
    out <- paste0('loom/', method, '_', covariate, '_', transform, '_', length(vfeatures), '_', rand, '_',  latent, '.loom')
	
	# add count to one cell if all are zero
	#vfeature_num <- length(vfeatures)
	#one0 <- vector(mode = 'numeric', length = vfeature_num)
	#one0[2] <- 1
	#if (sum(colSums(matrix)==0) > 0){
	#	matrix[,colSums(matrix) == 0] <- one0
	#}
	
	create(filename= out, 
           overwrite = TRUE,
           data = matrix, 
           cell.attrs = list(batch = seurat_obj@meta.data[,covariate],
                             batch_indices = seurat_obj@meta.data[,covariate] %>% 
                               as.factor() %>% 
                               as.numeric()))
    # connect to new loom file, then disconnect...otherwise python call gets borked for 
    # as we are connected into the file on create
    loom <- connect(out, mode = 'r')
    loom$close_all() 
    n_epochs = 30
    lr = 0.001 
    #use_batches = 'True'
    use_cuda = 'True'
    n_hidden = 128 
    n_latent = latent
    n_HVG = length(vfeatures) 
    
    scArches_command = paste('/data/mcgaugheyd/conda/envs/scArches/bin/python /home/mcgaugheyd/git/scEiaD/src/run_scArches.py',
                         out,
                         n_epochs,
                         lr,
                         use_cuda,
                         n_hidden,
                         n_latent,
                         n_HVG,
						 'FALSE')
	print(scArches_command)
    system(scArches_command)
    # import reduced dim (latent)
    latent_dims <- read.csv(paste0(out, '.csv'), header = TRUE)
	row.names(latent_dims) <- latent_dims[,1]
	latent_dims <- latent_dims[,-1]

    colnames(latent_dims) <- paste0("scArches_", 1:ncol(latent_dims))
    
    seurat_obj[["scArches"]] <- CreateDimReducObject(embeddings = latent_dims %>% as.matrix(), key = "scArches_", assay = DefaultAssay(seurat_obj))
	obj <- seurat_obj 
  } else if (grepl('scVI', method, ignore.case = TRUE)) { #method == 'scVI') {
    # scVI ----
    assay <- 'RNA'
    vfeatures <- grep('^MT-', seurat_obj@assays$RNA@var.features, invert =TRUE, value = TRUE)
    if (transform == 'counts'){
      matrix = seurat_obj@assays$RNA@counts[vfeatures, ]
    } else if (transform == 'SCT'){
      assay <- 'SCT'
      vfeatures <- grep('^MT-', seurat_obj@assays$SCT@var.features, invert =TRUE, value = TRUE)
      matrix = seurat_obj@assays$SCT@scale.data[vfeatures, ] %>% Matrix(., sparse = TRUE)
    } else {
      matrix = seurat_obj@assays$RNA@scale.data[vfeatures, ]
    }
	rand <- sample(1e7:9e7,1)
    out <- paste0('loom/', method, '_', covariate, '_', transform, '_', length(vfeatures), '_', rand, '_',  latent, '.loom')
	
	# add count to one cell if all are zero
	vfeature_num <- length(vfeatures)
	one0 <- vector(mode = 'numeric', length = vfeature_num)
	one0[2] <- 1
	if (sum(colSums(matrix)==0) > 0){
		matrix[,colSums(matrix) == 0] <- one0
	}
	seurat_obj@meta.data$SA = row.names(seurat__standard@meta.data) %>%  str_extract(., '(SRX|EGAF|ERS|SRS|iPSC_RPE_scRNA_)\\d+')	
	create(filename= out, 
           overwrite = TRUE,
           data = matrix, 
           cell.attrs = list(batch = seurat_obj@meta.data[,covariate],
						     sample_accession = seurat_obj@meta.data[,'SA'],
                             batch_indices = seurat_obj@meta.data[,covariate] %>% 
                               as.factor() %>% 
                               as.numeric()))
    # connect to new loom file, then disconnect...otherwise python call gets borked for 
    # as we are connected into the file on create
    loom <- connect(out, mode = 'r')
    loom$close_all() 
    #n_epochs = 5 # use 1e6/# cells of epochs
    lr = 0.001 
    #use_batches = 'True'
    use_cuda = 'True'
    n_hidden = 128 
    n_latent = latent
    n_layers = 2 
     
    scVI_command = paste(glue('{conda_dir}/envs/scvitools090/bin/./python {git_dir}/src/run_scVI2.py'),
                         out,
                         n_epochs,
                         lr,
                         use_cuda,
                         n_hidden,
                         n_latent,
                         n_layers,
						 'FALSE')
	if (method == 'scVIprojection') {
    	scVI_command = paste(glue('{conda_dir}/envs/scvitools090/bin/./python {git_dir}/src/run_scVI_projection.py'),
                         out,
                         n_epochs,
                         lr,
                         use_cuda,
                         n_hidden,
                         n_latent,
                         n_layers,
						 'FALSE')
	}
	if (method == 'scVIprojectionSO') {
    	scVI_command = paste(glue('{conda_dir}/envs/scvitools090/bin/./python {git_dir}/src/run_scVI_projection.py'),
                         out,
                         n_epochs,
                         lr,
                         use_cuda,
                         n_hidden,
                         n_latent,
                         n_layers,
						 'FALSE',
						 args[8],
						 args[9])
	}
    # run scVI     
	print(scVI_command) 
    system(scVI_command)
    # import reduced dim (latent)
	meta <- read_csv(paste0(out, '.meta.csv'))
	#ld <- read_csv('scVIlatent_MM-MF_projected_to_HS.csv')
		
	ld <- read_csv(paste0(out, '.latent.csv'))
	metald <- cbind(meta, ld[,2:ncol(ld)])
	ld_ordered <- seurat__standard@meta.data %>% as_tibble(rownames = 'Barcode') %>% left_join(metald %>% mutate(Barcode = gsub('-1$|-0$','', X1)), by = c('Barcode'))
	scVI_cols <- grep('^\\d+$', colnames(ld_ordered))
	ld_ordered <- ld_ordered[,scVI_cols] 
	colnames(ld_ordered) <- paste0("scVI_", 1:ncol(ld_ordered))
	ld_ordered <- ld_ordered %>% as.matrix()
	row.names(ld_ordered) <- colnames(seurat__standard)
	seurat_obj[["scVI"]] <- CreateDimReducObject(embeddings = ld_ordered, key = "scVI_", assay = DefaultAssay(seurat_obj))	
#seurat_obj <- SetAssayData(object = seurat_obj, slot = 'scale.data', new.data = normalized_values)
	#save(normalized_values, file = gsub('.seuratV3.Rdata', '.scVI_scaled.Rdata', args[6]), compress = FALSE)
	# system(paste0('rm ', out, '.normalized.csv'))
	obj <- seurat_obj 
    
  } else if (method == 'harmony'){
    ## uses one seurat obj (give covariate in meta.data to group.by.vars)
    if (transform != 'SCT'){
      obj <- RunHarmony(seurat_obj, group.by.vars = covariate,
                        max.iter.harmony = 5, 
                        epsilon.harmony = -Inf)
    } else {
      obj <- RunHarmony(seurat_obj, group.by.vars = covariate,
                        max.iter.harmony = 5, 
                        epsilon.harmony = -Inf,
                        assay.use = 'SCT')
    }
  } else if (method == 'liger'){
    ## like harmony above, give one seurat obj
    ## NMF requires POSITIVE values
    ## so, UNLIKE, the tutorial set the lowest value to 0.1 in the matrix
    var_genes <- grep('^MT-', seurat_obj@assays$RNA@var.features, value = TRUE, invert = TRUE)
    obj <- seurat_obj
    # identify batches wiht really low cell coutns (<100) to exclude
    obj@meta.data$split_by <- obj@meta.data[,covariate]
    #splits_to_remove <- obj@meta.data %>% 
    #  dplyr::group_by(split_by)%>% 
    #  summarise(Count = n())  %>% 
    #  filter(Count < 100) %>% 
    #  pull(split_by) 
    #if (length(splits_to_remove >= 1)) { 
    #  obj <- subset(obj, subset = split_by %in% splits_to_remove, invert = TRUE)
    #}
    obj <-  ScaleData(seurat_obj, split.by = covariate, do.center = FALSE)
    #obj@assays$RNA@scale.data <- obj@assays$RNA@scale.data - min(obj@assays$RNA@scale.data) + 0.1 # <- yeah this is hacky but I think OK...
    # ...the alternative would be to re-run from scratch with raw counts, which would mean 
    # liger would be getting differently scaled values
    # than the other methods...
    obj <- RunOptimizeALS(obj, 
                          k = latent, 
                          lambda = 5, 
                          split.by = covariate)
    obj <- RunQuantileAlignSNF(obj, split.by = covariate)
    # finally re-do scaledata in case I use it in the future...I prob won't be expecting
    # it to not be centered and with no neg values....
    #obj <- ScaleData(obj,  
    #                 features = var_genes,
    #                 do.center = TRUE,
    #                 do.scale = TRUE,
    #                 vars.to.regress = c("nCount_RNA", "nFeature_RNA", "percent.mt"))
  } else if (method == 'scanorama'){
    # scanorama ----
    # scanorama can return "integrated" and/or "corrected" data
    # authors say that the "integrated" data is a low-dimension (100) representation
    # of the integration, which is INTENDED FOR PCA/tSNE/UMAP!!!
    # the corrected data returns all of the sample x gene matrix with batch
    # corrected values
    if (transform != 'SCT'){
      assay <- 'RNA'
    } else {assay <- 'SCT'}
    d <- list() # d is lists of scaled Data
    g <- list() # g is lists of gene names for each matrix in d
    seurat_list <- SplitObject(seurat_obj, split.by = covariate)
    # build name-less list of expression values
    # and name-less list of gene names
    # if you give a list with names then reticulate / python integration explodes
    # well actully just the name are given to python...which took me
    # forever to figure out. stupid annoying. 
    for (i in seq(1, length(seurat_list))){
      # print(i);
      var_genes <- grep('^MT-', seurat_list[[i]]@assays[[assay]]@var.features, value = TRUE, invert = TRUE)
      d[[i]] <- t((seurat_list[[i]]@assays[[assay]]@scale.data[var_genes,])) %>% as.matrix(); 
      d[[i]][is.na(d[[i]])] <- 0; 
      g[[i]] <- colnames(d[[i]]) 
    }
    print('Run scanorama')
    integrated.corrected.data <- 
      scanorama$correct(d, g , return_dimred=TRUE, return_dense=TRUE)
    print('Scanorama done')
    # get the cell names back in as they aren't returned for some reason (reticulate?)
    for (i in seq(1, length(d))){
      # first is in the integrated data (dim reduced for UMAP, etc)
      row.names(integrated.corrected.data[[1]][[i]]) <- row.names(d[[i]])
    }
    
    # glue reduced matrix values into seurat for later UMAP, etc
    scanorama_mnn <- Reduce(rbind, integrated.corrected.data[[1]])
    colnames(scanorama_mnn) <- paste0("scanorama_", 1:ncol(scanorama_mnn))
    obj <- seurat_obj
    obj[["scanorama"]] <- CreateDimReducObject(embeddings = scanorama_mnn, key = "scanorama_", assay = DefaultAssay(obj))
    
  } else if (method == 'combat') {
    if (transform != 'SCT'){
      assay <- 'RNA'
    } else {assay <- 'SCT'}
    # gene by sample for ComBat
    obj <- seurat_obj
    var_genes <- grep('^MT-', obj@assays[[assay]]@var.features, value = TRUE, invert = TRUE)
    matrix <- obj@assays[[assay]]@scale.data[var_genes,]
    cor_data = ComBat(matrix, obj@meta.data[, covariate], prior.plots=FALSE, par.prior=TRUE)
    obj <- SetAssayData(obj, slot = 'scale.data', cor_data)
    obj <- RunPCA(obj, npcs = 100)
  } else {
	print(method)
    print('Supply either CCA, fastMNN, harmony, liger, scanorama, or scVI as a method')
    NULL
  }
  obj
}

if (transform != 'SCT' & grepl('scVI', method)){
  integrated_obj <- run_integration(seurat__standard, method, covariate, transform, latent = latent, n_epochs = as.integer(args[7]))
} else if (transform != 'SCT' & method != 'none'){
  integrated_obj <- run_integration(seurat__standard, method, covariate, transform, latent = latent)
} else if (transform == 'SCT' & method == 'CCA') {
  integrated_obj <- run_integration(seurat__SCT, 'CCA', covariate, transform = 'SCT', latent = latent)
} else if (transform == 'SCT' & method != 'none') {
  seurat_list <- seurat__SCT$seurat_list
  if (length(seurat_list) > 1){
    merged <- merge(x = seurat_list[[1]], y = seurat_list[2:length(x = seurat_list)])
  } else {merged <- seurat_list[[1]]}
  merged@assays$SCT@var.features <- seurat__SCT$study_data_features
  DefaultAssay(merged) <- 'SCT'
  merged <- RunPCA(merged, npcs = 100)
  integrated_obj <- run_integration(merged, method, covariate, transform = 'SCT', latent = latent)
} else if (transform != 'SCT' & method == 'none'){
  integrated_obj <- seurat__standard
} else if (transform == 'SCT' & method == 'none'){
  seurat_list <- seurat__SCT$seurat_list
  if (length(seurat_list) > 1){
    merged <- merge(x = seurat_list[[1]], y = seurat_list[2:length(x = seurat_list)])
  } else {merged <- seurat_list[[1]]}
  merged@assays$SCT@var.features <- seurat__SCT$study_data_features
  DefaultAssay(merged) <- 'SCT'
  merged <- RunPCA(merged, npcs = 100)
  integrated_obj <- merged
}
save(integrated_obj, file = args[6], compress = FALSE)
```
